# YASS: Yet Another Spike Sorter applied to large-scale multi-electrode array recordings in primate retina

**DOI:** 10.1101/2020.03.18.997924

**Authors:** JinHyung Lee, Catalin Mitelut, Hooshmand Shokri, Ian Kinsella, Nishchal Dethe, Shenghao Wu, Kevin Li, Eduardo Blancas Reyes, Denis Turcu, Eleanor Batty, Young Joon Kim, Nora Brackbill, Alexandra Kling, Georges Goetz, E.J. Chichilnisky, David Carlson, Liam Paninski

## Abstract

Spike sorting is a critical first step in extracting neural signals from large-scale multi-electrode array (MEA) data. This manuscript presents several new techniques that make MEA spike sorting more robust and accurate. Our pipeline is based on an efficient multi-stage “triage-then-cluster-then-pursuit” approach that initially extracts only clean, high-quality waveforms from the electrophysiological time series by temporarily skipping noisy or “collided” events (representing two neurons firing synchronously). This is accomplished by developing a neural network detection and denoising method followed by efficient outlier triaging. The denoised spike waveforms are then used to infer the set of spike templates through nonparametric Bayesian clustering. We use a divide-and-conquer strategy to parallelize this clustering step. Finally, we recover collided waveforms with matching-pursuit deconvolution techniques, and perform further split-and-merge steps to estimate additional templates from the pool of recovered waveforms. We apply the new pipeline to data recorded in the primate retina, where high firing rates and highly-overlapping axonal units provide a challenging testbed for the deconvolution approach; in addition, the well-defined mosaic structure of receptive fields in this preparation provides a useful quality check on any spike sorting pipeline. We show that our pipeline improves on the state-of-the-art in spike sorting (and outperforms manual sorting) on both real and semi-simulated MEA data with > 500 electrodes; open source code can be found at https://github.com/paninski-lab/yass.

## 1 Introduction

The analysis of large-scale multineuronal spike train data is crucial for current and future neuroscience research. These analyses are predicated on the existence of reliable and reproducible methods that feasibly scale to ever-increasing data acquisition rates. A standard approach for collecting these data is to use dense multi-electrode array (MEA) recordings followed by “spike sorting” algorithms to turn the obtained raw electrical signals into spike trains.

Currently widely-employed MEAs enable recordings at a scale of ~ 10^4^ electrodes (Jun et al., 2017a), but efforts are underway to increase this number to 10^6^ electrodes^‡^. At this scale any manual processing of the obtained data is infeasible. Therefore, automatic spike sorting for dense MEAs has enjoyed significant recent attention (Franke et al., Carlson et al., 2014, Kadir et al., Ekanadham et al., Muthmann et al., 2015, Pachitariu et al., 2016, Yger et al., 2018, Hilgen et al., 2017, Jun et al., 2017b). Despite these efforts, spike sorting remains a critical computational bottleneck in the scientific pipeline when using dense MEAs, due both to the high computational cost of the algorithms and the human time spent on manual postprocessing.

To accelerate progress on this critical scientific problem, we propose methodology guided by several main principles. First, robustness should be critical, since hand-tuning and post-processing is not feasible at the large scales found in current and future MEA datasets. Second, prior information should be leveraged as much as possible; we share information across neurons, electrodes, and experiments in order to extract information from the MEA datastream as efficiently as possible. Finally, scalability must be a key consideration. To feasibly process the oncoming data deluge, we use parallel, scalable algorithms based on efficient data summarizations wherever possible and focus computational power on the “hard cases,” using cheap fast methods to handle easy cases.

To evaluate the resulting pipeline, we focus here on MEA data collected from the primate retina. This preparation is a useful spike sorting testbed for several important reasons. First, the two-dimensional MEA used here matches the approximately two-dimensional substrate of the retinal ganglion layer. Second, receptive fields of well-characterized retinal ganglion cell (RGC) types (e.g., ON parasols, OFF midgets, etc.) are known to approximately tile the visual field, providing useful side information for scoring different spike sorting pipelines. Third, many RGCs have moderately high firing rates and often have significant axonal projections that overlap with each other spatially on the MEA, making it challenging to demix spikes that overlap spatially and temporally from different RGCs*.

We will first outline the methodology that forms the core of our pipeline in Section 2.1, then provide details of each module in the following subsections, and finally demonstrate the improvements in performance on 512-electrode primate retina MEA recordings in Section 3.

## 2 Methods

### 2.1 Overview

Our overall strategy can be considered a hybrid of a sparse deconvolution approach (specifically, an extension of the matching pursuit method used in (Pachitariu et al., 2016)) and a classical clustering approach, generalized and adapted to the large dense MEA setting. Our guiding philosophy is that it is essential to properly handle “collisions” between simultaneous spikes (Pillow et al., Ekanadham et al., Pachitariu et al., 2016), since collisions distort the extracted feature space and hinder clustering. (These collisions are quite prevalent in primate retinal data, as illustrated in Figure 1.) Sparse deconvolution approaches can potentially handle these collisions, but require good initializations of each neuron’s “template” (average spike shape). Our approach therefore “triages” collided or noisy waveforms in the initial template estimation step, excluding these corrupted spikes from the feature extraction and clustering stages, and deferring recovery of such spikes to later deconvolution stages. This leads to significantly improved template estimates, and therefore more robust and accurate overall results.

**Figure 1:**
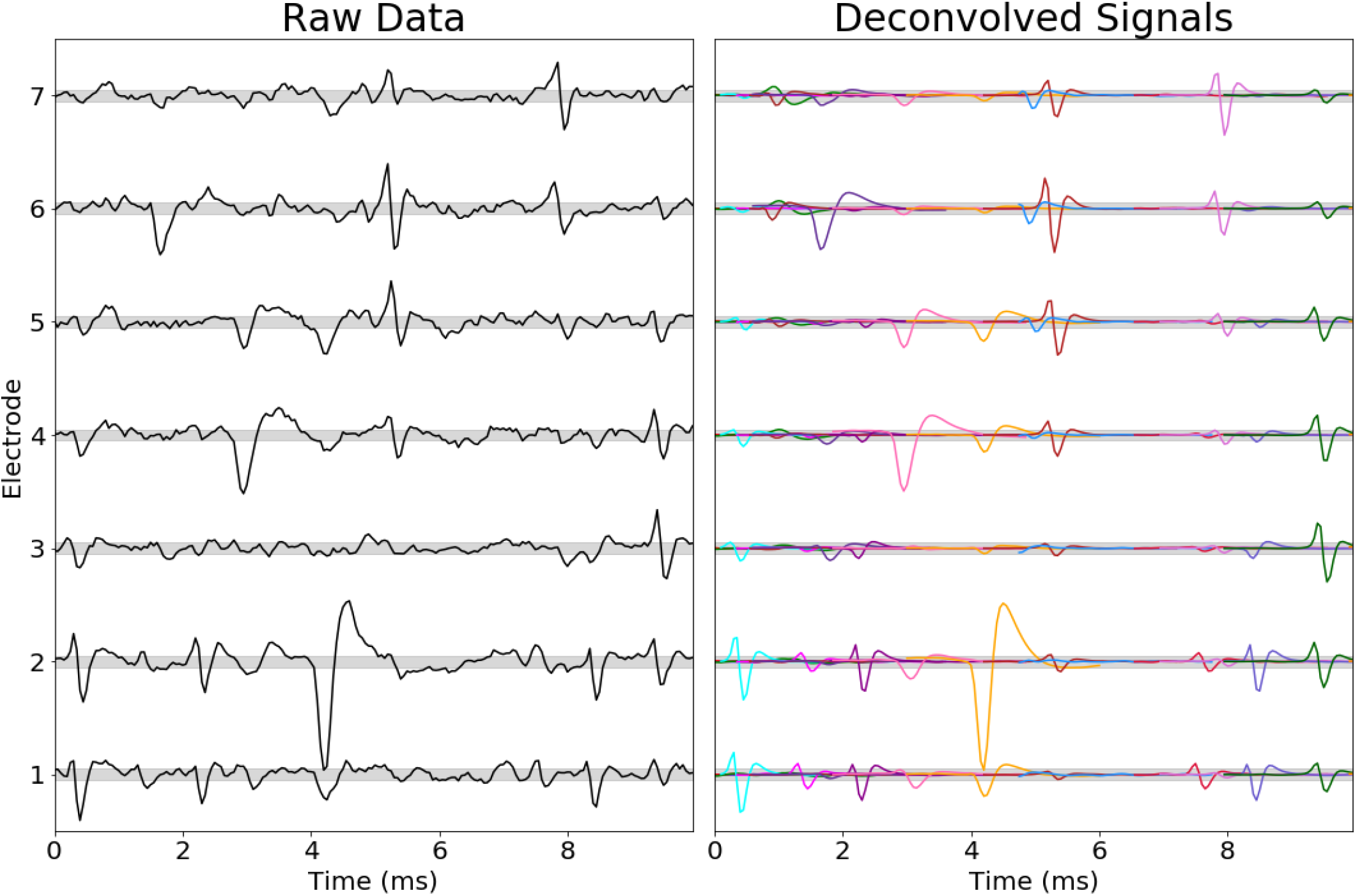
High per-electrode firing rates and the prevalence of collisions in primate retina data. (Left) voltage signal on 7 neighboring channels during a randomly selected 10 ms interval; (right) spike shapes and times identified and sorted by YASS (each unit is assigned a random color). The gray shade (here and in the following figures) represents ±1 estimated noise scale (estimated as the median absolute deviation of the raw data on each channel, divided by 0.674). Note that some of the spikes visible on these electrodes are small but are larger on other electrodes (not shown); all units shown here have templates with peak-to-peak amplitude > 4.5× the estimated noise scale on at least one electrode. The total firing rate (summed over all visible units) on each channel here is in the hundreds of Hz range, leading to frequent collisions.

Guided by this overall philosophy, our approach is a modular, multistage pipeline that includes the following main stages: (*i*) detection, denoising, and de-duplication of spike waveforms, (*ii*) division of the waveforms into distinct subsets for parallel processing, (*iii*) clustering the waveforms using adaptive featurization while triaging outliers and collided waveforms to obtain single neuron templates, and finally (*iv*) inferring missed and collided spikes via a sparse deconvolution step. Pseudocode for the flow of the pipeline can be found in Algorithm 1. Below we provide a summary overview of each stage with the following subsections describing in detail each of the modules.

We start by band-pass filtering and standardizing the voltage signals using standard methods. For completeness, these steps are described in section 2.2.

The next stage is event or spike *detection* (section 2.3), implemented by a convolutional neural network (NN). The network is trained using simulated data that is designed to emulate the correlated noise and spike shapes observed in real data; no hand-labeling of individual spikes is needed.

Detected spike waveforms are then *denoised* using a network that is trained using a similar approach as the detection network (section 2.4). The denoising process also serves to suppress corruptions of spike shapes due to collisions^†^.

The detection NN operates locally in space and time; spatially-extended axonal spikes may be detected in multiple locations. Therefore, we implement a spike de-duplication step before further processing (section 2.5).

Next we upsample and align spikes to the mean spike shape on each channel (section 2.6).

A preliminary clustering step follows that uses denoised waveforms (section 2.7). Clustering this large and high-dimensional data is challenging; several main ideas are critical to make this clustering step tractable. First, we use a coarse-to-fine divide-and-conquer approach: instead of trying to run a single clustering algorithm on the full dataset, we iteratively split the data into smaller subsets (starting by examining each denoised waveform on its main channel — i.e., the electrode on which the spike is biggest — and then performing more splits on secondary channels) before clustering these smaller subsets of data. Second, we do not attempt to cluster the full high-dimensional waveforms; instead, we perform a featurization within each subset (adapting the chosen features to each subset) and cluster in the estimated low-dimensional feature space. Third, we triage outliers (caused mostly by collisions) that would otherwise lead to clustering errors. Finally, we apply a nonparametric Bayesian approach (similar to (Miller and Harrison, 2018)) with stochastic variational split and merge methods to efficiently explore each clustering space (Hughes and Sudderth) and automatically estimate the number of clusters in each subset.

#### Algorithm 1 Pseudocode for the complete proposed pipeline

~~~
 1: Input: raw recording
 2:
 3: **function** Preprocess (sec. 2.2)(raw recording)
 4:    **return** standardized recording
 5:
 6: **function** Spike Extraction(first batch of standardized recording)
 7:    Detection (sec. 2.3)
 8:    Denoising (sec. 2.4)
 9:    De-duplication (sec. 2.5)
10:    **return** spike waveforms
11:
12: **function** Initial Sort(spike waveforms)
13:    Alignment (sec. 2.6)
14:    Clustering (sec. 2.7), Cluster Post-process (sec. 2.8)
15:    **return** templates
16:
17: **function** Initial Deconvolution (templates, first batch of standardized recording)
18:    Deconvolution (sec. 2.9)
19:    **return** spike train
20:
21: **function** Re-sort(spike train, templates, first batch of standardized recording)
22:    Alignment (sec. 2.6), Denoising (sec. 2.4)
23:    Post-deconvolution Reclustering (sec. 2.10.2), Cluster Post-process (sec. 2.8)
24:    Deconvolution (sec. 2.9), Post-deconvolution Merge (sec. 2.10.3)
25:    Soft Assignment (sec. 2.12)
26:    **return** templates, spike train
27:
28: **function** Final Deconvolution(templates, remaining batches of standardized recording)
29:    Deconvolution with Drift Handling (sec. 2.11)
30:    Soft Assignment (sec. 2.12)
31:    **return** time-varying templates, spike train, soft assignment
32:
33: Output: time-varying templates, spike train, soft assignment
~~~

In practice, the divide-and-conquer approach described above can oversplit some units, and sometimes forms small clusters that are dominated by collisions, instead of well-isolated single spikes. We developed some simple post-processing steps to handle both of these issues (section 2.8).

The cluster means or neuron templates yielded by the clustering stage are then used to infer the presence of all spikes (whether isolated or collided) across the entire dataset, using a deconvolution algorithm (section 2.9). We use a matching pursuit approach similar to that employed in Kilosort (Pachitariu et al., 2016), but with a few important modifications, including super-resolution temporal spike alignment and an iterative coordinate-descent (ICD) step that helps correct some of the errors made by the initial greedy matching pursuit pass.

Much of the variability in the shape of spikes in primate retinal recordings is due to collisions with other spikes. After the deconvolution step, for each spike, we can estimate the contributions of all the other inferred spikes, and regress these contributions away to effectively “clean” each observed spike. We then run additional split/merge steps on these post-deconvolution cleaned waveforms to update the templates (section 2.10).

To capture template scale changes during longer recordings, we implement a simple exponentially-weighted update algorithm, partially similar to the basic approach used in Kilosort2 (Pachitariu, 2019) (section 2.11).

Finally, low signal-to-noise-ratio (SNR) spikes from different cells can be very similar to each other; to properly handle uncertainty in the assignments of these spikes, we use a simple probabilistic model of the noise in the deconvolution step to compute soft assignments (section 2.12).

YASS is written in Python and CUDA; open source code can be found at https://github.com/paninski-lab/yass. The preprocessing, spike detection, denoising, and deconvolution steps are performed on temporal minibatches of data (parallelized over multiple GPUs, if available) in order to scale to large datasets; the other pipeline stages are parallelized over multiple CPU cores and operate on significantly reduced data representations to limit memory usage. The pipeline has been tested on PCs, multi-processor workstations, and on Amazon Web Services (AWS).

### 2.2 Preprocessing

The recording is first band-pass filtered using a Butterworth filter (300Hz-2KHz for the data analyzed here) then standardized by the Median Absolute Deviation (MAD).

### 2.3 Detection

Next we detect putative spike events. The optimal detection of spikes with unknown shapes and sizes, in the presence of collisions and spatiotemporally correlated noise, is a challenging statistical problem; simple thresholding is computationally straightforward but clearly statistically suboptimal. (Classical signal detection theory tells us that linear filtering followed by thresholding is the optimal approach for detecting a known signal shape in Gaussian noise — but here the signal shape is unknown and the “noise” is dominated by collisions and is highly non-Gaussian.) Typically an experimentalist will have very strong priors about what a “good spike” will look like, and a statistically-optimal approach will take advantage of this strong prior information — but it is not obvious how to encode these priors in a computationally-tractable detection algorithm.

Neural networks (NNs) have emerged as the current state-of-the-art method for translating large labeled datasets into highly-accurate classifiers or denoisers. However, we do not want to spend time hand-labeling many individual spikes here. Instead, we begin with a reasonable generative model that outputs short, small spatiotemporal snippets of voltage data, then sample at will from this generative model to provide as many training minibatches to the neural network as desired. When training converges, we have a trained NN that (locally) optimizes the classification accuracy under the generative model. Thus this approach is a method that translates the priors made explicit in a generative model into an optimal classifier that implicitly encodes these priors. See e.g. (Parthasarathy et al., 2017, Yoon et al., 2017, Weigert et al., 2018, Sun and Paninski, 2018) for previous applications of similar ideas to image denoising and decoding problems.

Thus, given the availability of standard NN training routines, we have shifted most of the effort here onto the definition of the generative model. We use a straightforward convolutional model to generate the required spatiotemporal voltage snippets: we sample spike shapes from some library, scale these shapes randomly, choose random times for the spikes, and linearly sum the result along with a sample from a stationary spatiotemporally-correlated noise process. We have found that a Gaussian process model for the noise suffices to obtain strong classification results; for the spike shape library, we simply collect templates from earlier successful sorts from the same preparation. Thus only very modest user input is required here, in the selection of “good” spike templates. See Figure 2 for an illustration, and the appendix for full details on the generative model and NN training.

**Figure 2:**
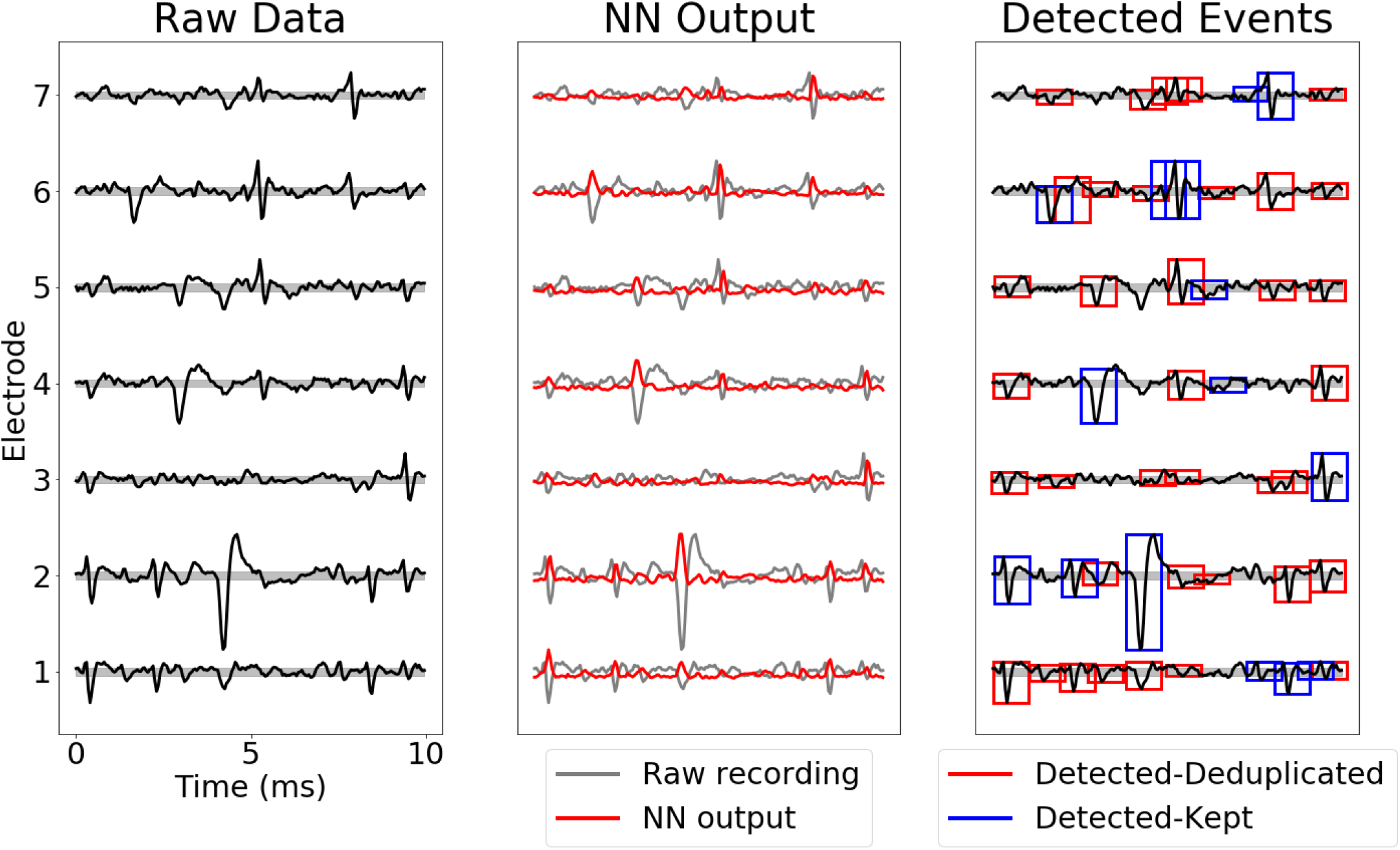
Illustration of detection and de-duplication steps. (**Left**) Raw recording on seven neighboring electrodes; (**Middle**) Neural Network (NN) output (red) overlaid on top of raw recording. (**Right**) detected and de-duplicated spikes. Red (blue) boxes denote spikes that are removed (retained) during the de-duplication step.

### 2.4 Denoising

Single detected spikes often appear quite noisy; in the primate retinal preparation, much of this heavy-tailed “noise” is due to collisions with other spikes (recall Figure 1). It is useful to suppress this “noise” before further processing; the goal here is to input a noisy, potentially collision-corrupted spike and output a “denoised” version of the spike. As in the detection context discussed above, simple linear denoising (e.g., projection onto a subspace inferred by principal or independent components analysis) is suboptimal here, due to the strongly non-Gaussian nature of collisions. To proceed, we use the same trick that we used for detection: we use the same generative model to create an arbitrarily large training set and then train a NN to solve this denoising task, i.e., take corrupted noisy spikes and output the original uncorrupted spike waveform. See Figure 3 for an illustration, and the appendix for full architecture and training details.

**Figure 3:**
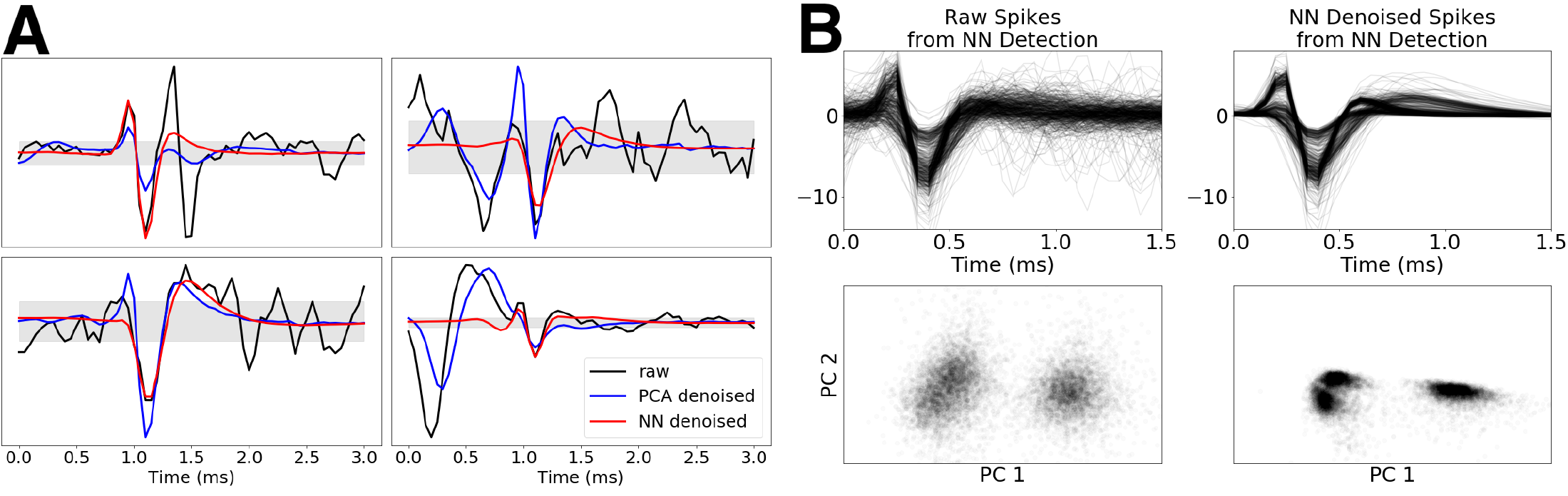
Illustration of denoising step. **(A)**: Four examples of raw detected spikes (black) and corresponding NN-denoised spikes (red) and PCA projections (blue). Note that the NN successfully suppresses both noise and contributions from other collided spikes, while the PCA projection is significantly more sensitive to collisions. **(B)**: (Top) raw detected spikes collected on a single electrode (left) vs. the same spikes after denoising (right). (Bottom): PCA projections corresponding to the raw (left) vs denoised spikes (right). Note that many of the outliers are no longer visible in the PCA projections, and the three clusters are much more easily distinguishable by eye.

### 2.5 De-duplication of detected events

Our detection NN operates locally on just a few neighboring electrodes to detect spikes. In the datasets analyzed here, single spikes are often visible over hundreds of electrodes (see Figure 5 below for examples of some spatially-extended spike templates). Therefore single spikes may be detected simultaneously on many different electrodes, and before further processing we need to at least partially de-duplicate these multiple detected events.

**Figure 4:**
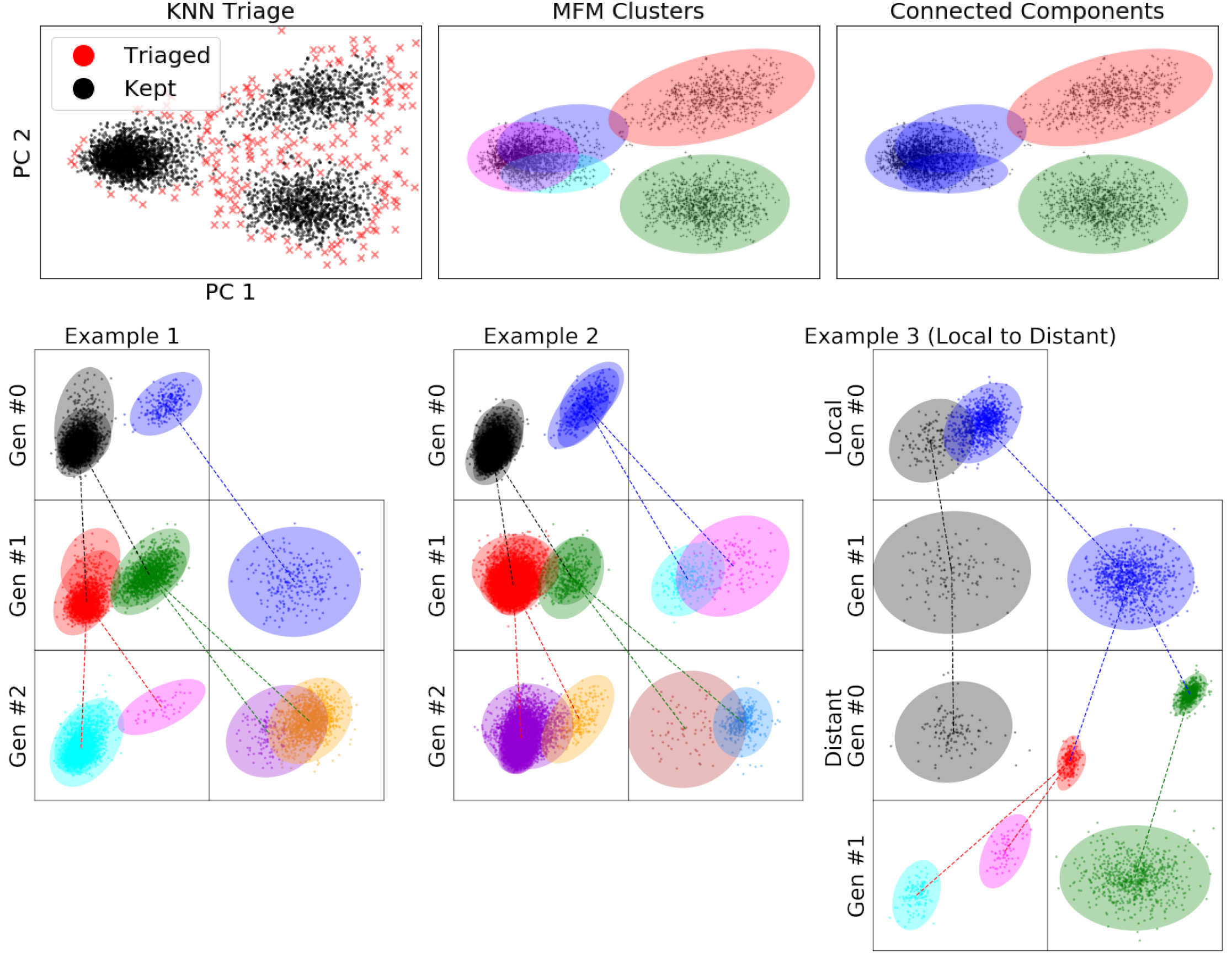
Divide-and-conquer clustering with triaging and iterative re-featurization. (Top left): Outlier triaging. Top middle: mixture-of-finite-mixture model (MFM) clustering of datapoints remaining after triage step. Each color represents a single cluster returned by MFM run in a higher-dimensional feature space. Top right: grouping of clusters with substantial overlap using connected components. (Bottom row): Three examples illustrating the benefits of iterative re-featurization. Each box represents a single clustering iteration. Ellipses represent MFM clusters and colors represent connected components (CCs); while the actual featurization is five-dimensional, for visualization the five-dimensional vectors are projected on the two-dimensional feature space that best separates the clusters (via linear discriminant analysis). In each “generation” we perform a triage, then an MFM clustering, then a connected-components analysis (as in the top row); then we re-featurize the data within each CC and iterate in the next generation. Colored lines indicate the flow of data in this procedure: a CC in one generation might generate multiple CC in the following generation, if the new featurization is informative enough to split clusters that were not separable in the previous generation. Indeed, as shown in Examples 1 and 2, many CCs can be more clearly split in the next generation by adaptive re-featurization. Example 3 illustrates the necessity of features from distant electrodes. In the first two rows, clustering uses features from local electrodes only, and the last two rows utilize features from distant electrodes. Although MFM returns a single blue cluster in generation 1 of local clustering, re-featurizing using distant features shows two clearly separable red and green CCs in generation 0 using distant features. See also Figure 5 below for an illustration of some units that are only separable via distant features.

**Figure 5:**
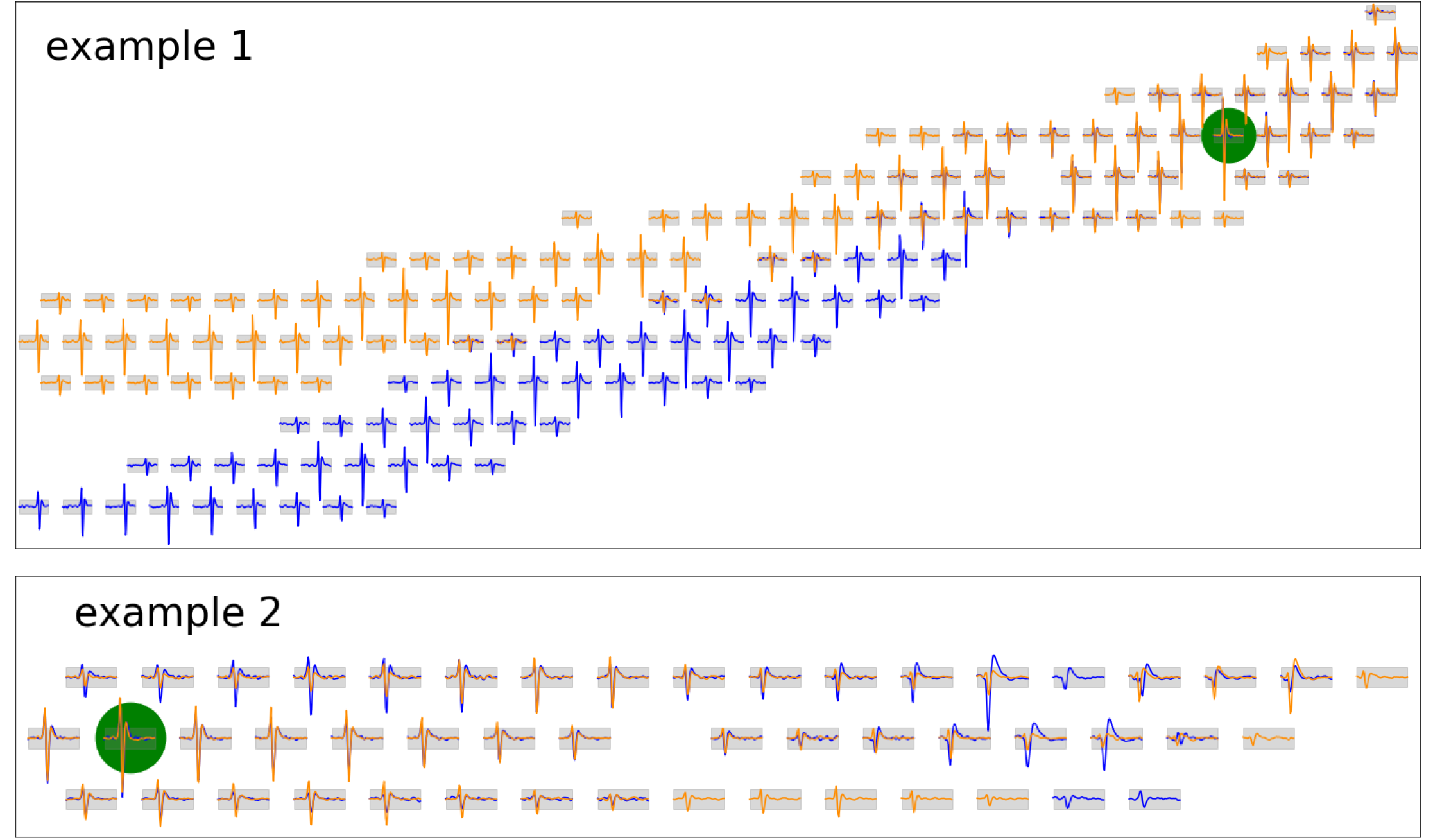
Examples illustrating the need for distant-electrode featurization. Two example pairs of units whose templates look locally similar but can be easily separated if we use information from all electrodes in the clustering process. In each example, the green dot represents the (shared) primary channel of the two templates indicated in blue and orange. Only “active” channels (where the template height is above a threshold) are shown.

Full de-duplication of single spikes, without knowledge of the complete set of spike templates, is a statistically-and computationally-challenging problem that we do not attempt to solve. Instead, for each detected spike, we simply search locally in a spatiotemporal neighborhood for other smaller synchronous events, and drop any smaller events that we find. (Specifically, we use a window of 0.25*ms* and 150*μm* for this search, and compare the size of denoised events in terms of their peak-to-peak spike heights.) This de-duplication is fast but decidedly imperfect: it will kill duplicate detected events that come from the same cell but are smaller than the spike on the cell’s “primary” channel (where the spike tends to be largest), as desired — but it will fail to de-duplicate very spatially extended spikes that extend outside of the spatial de-duplication window (false positives), and it will also kill spikes from smaller cells that happen to fire synchronously with the bigger cell (false negatives). We will correct the first issue with a template-deduplication step (section 2.8.2), and handle the second issue in the deconvolution step (section 2.9).

### 2.6 Alignment

Next we temporally align the de-duplicated waveforms. In practice, we find that super-resolution alignment (i.e., upsampling and then aligning waveforms) further helps reduce spike waveform variability, leading to better downstream clustering. Specifically, we use spline interpolation to upsample the raw waveform of each spike on its primary channel (to 5× the acquisition rate), determine optimal shifts for each spike, and then use linear interpolation to apply this alignment to all secondary channels. The optimal shift is determined by maximizing the dot product of a shifted waveform with the average of all waveforms (separated by their primary channel).

### 2.7 Coarse-to-fine divide-and-conquer clustering with triaging and iterative re-featurization

At this point, for each electrode *i*, we have an array of detected spikes which are denoised, (partially) de-duplicated, aligned, and are largest on electrode *i* (i.e., we have grouped each spike by its primary electrode). Now, finally, we turn to clustering each of these datasets in parallel.

This clustering problem is challenging in several key respects. First, even after grouping detected spikes by their primary channel, the data remains very large: in the datasets considered here, we have to handle thousands of spikes per channel per minute, with tens of thousands of potential features per spike as some axonal spikes can appear on up to 100 channels (or more in some cases), with each channel containing ~ 60 timepoints. Second, as emphasized above, the variability here is highly non-Gaussian, due largely to collisions; the NN denoiser suppresses many but not all collisions. Third, the number of clusters per channel is *a priori* unknown. Finally, while most cells can be separated using only local features from electrodes near the primary electrode, for some cells we do need to include features from more distant electrodes to achieve good cluster separation; see Figure 5 below.

To address these challenges, we combine several key ideas:

**Divide-and-conquer**. The first step is to try to divide the dataset into smaller subsets before applying any expensive clustering algorithms (Swindale and Spacek (2014)). This “divide-and-conquer” approach is critical for both statistical and computational reasons. Naive clustering has a computational cost that scales superlinearly with both the number of datapoints and the dimensionality of the feature space. If we can split the data into subsets containing clearly different units, this would reduce both the number of datapoints per clustering call and the feature dimensionality needed to split the clusters; in turn, this dimensionality reduction can reduce overfitting given limited data. Finally, splitting the data improves parallelism; running more smaller jobs tends to be more scalable than running fewer bigger jobs.

**Coarse-to-fine**. Of course, all the above points are irrelevant if we can’t easily find ways to split the data. Luckily, for our problem, a coarse-to-fine strategy works quite well. We begin at the coarsest level with an extremely simple, one-dimensional featurization of the data: the size of the spike on the primary channel (as measured by the peak-to-peak height of the denoised spike), and perform a quick clustering using this feature, splitting the clearly distinct clusters into separate groups and leaving overlapping clusters in the same group. Next we incorporate “local” features (from neighbor electrodes to the primary electrode only) and perform further splits. Finally, at the “finest” level of the hierarchy, we incorporate features from more distant electrodes to perform a final set of splits.

**Iterative re-featurization**. The optimal featurization of the data might vary strongly from one subset of data to another; e.g., some clusters may be highly distinguishable based on the primary electrode, while for other clusters we might need to examine the secondary electrodes to achieve accurate splits. Therefore we do not start by reducing the dimensionality of the data (and then apply the same low-dimensional featurization to all the following splits); instead, we re-featurize after each split, i.e., compute a new dimensionality reduction of the data within each split subset. Figure 4 illustrates this process.

**Outlier triaging**. After dimensionality reduction, many outliers (largely due to collisions) are visible in the feature space. Instead of trying to assign these outliers to clusters (which leads in practice to unstable cluster shapes and over-splitting), we triage outliers — i.e., exclude them from the clustering step. (We will recover collision-corrupted spikes later in the pipeline during the deconvolution step described in section 2.9.) See the top-left panel of Figure 4 to see this triaging step at work.

**Nonparametric Bayesian clustering**. Finally, once we have chosen a good featurization and triaged outliers, we can choose a clustering algorithm. We suspect that there are many clustering algorithms that could work well at this stage; we have chosen a nonparametric Bayesian approach based on a “mixture of finite mixtures” (MFM) model (Miller and Harrison, 2018) that provides good results. Critically, the resulting algorithm outputs probabilistic estimates of cluster membership for each data point and is able to choose a reasonable number of clusters in a data-driven manner, without user input.

We discuss several of these issues in more detail in the following subsections; as usual, full mathematical and algorithmic details are provided in the appendix.

#### 2.7.1 Featurization for clustering using local or distant electrodes

During the “local” clustering phase of the coarse-to-fine approach, we start with denoised waveforms from the primary and nearest-neighbor electrodes. For hexagonal MEAs commonly used for retinal recordings, we considered 7 channels (i.e. the spike’s primary channel and the 6 neighboring channels), with 60 time points from each channel (3 ms for 20 kHz sampling rate), resulting in a vector with about 400 dimensions. We next randomly select up to 10,000 spikes on each channel to limit the total memory consumption.

After splitting clusters using local features in the previous step, in the “distant” clustering phase we restrict attention to features from more distant electrodes. Here we need to restrict attention to electrodes which might provide useful information for additional splits — trying to include all time points from all electrodes in the feature vector would lead to poor results due to overfitting. First we drop any “inactive” electrodes for which the mean voltage for this group of spikes is below a threshold (about 1 standardized voltage unit, i.e., about the noise scale). Second, to find electrodes and time points where there is some significant variability that may be due to the presence of multiple clusters, we compute the MAD of the waveforms on these electrodes, select the voltages at any time points where these values are greater than a threshold (again on the order of one standardized voltage unit)^‡^, and append these voltages into the feature vector. The resulting feature dimensionality ranges from 5 to 80.

Finally, we take the feature vectors computed above and then apply PCA with 5 components to reduce the dimensionality.

#### 2.7.2 Triaging

We use a simple, fast K-nearest-neighbors (K-NN) approach to detect outliers, following Knox and Ng (1998). When the feature dimensionality is small (recall here that our feature dimensionality is 5, following principal components projection), a kd-tree can find neighbors in 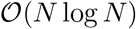 average time given a set with *N* elements. After computing the nearest neighbor graph we compute the average distance of each point to its five nearest neighbors (this can be thought of as a crude estimate of the local inverse density at each point), and triage out the top 1% datapoints with the largest average distance. This method effectively identifies outliers (Figure 4, top left panel).

#### 2.7.3 Cluster stability and connected component grouping

After running the nonparametric Bayesian (MFM) clustering algorithm (described in full detail in the appendix) we want to form groups of similar clusters and then re-featurize and re-cluster within these groups.

We begin by computing the “stability” of each cluster *i*, defined as the average of the assignment probabilities *p_ij_* that a point *j* is assigned to cluster *i* over all points *j* assigned with high probability to cluster *i*. Clusters that have a stability >0.90 are considered well isolated; we separate these clusters out and re-featurize and re-cluster their assigned spikes. This method is repeated recursively until MFM returns a single cluster; this is the termination point of the recursion.

In cases where none of the individual clusters returned by MFM has a stability >0.90, we group the pair of MFM-clusters that are closest (in Mahalanobis distance) and then recompute the stability metric on this group. We continue to do this until there is at least one group with >0.90 stability. See Figure 4, top, for an illustration of the resulting connected components.

### 2.8 Cluster post-processing

The clustering stage yields neuron templates, i.e. neuron shapes computed by averaging the aligned waveforms assigned to each cluster. Due to collisions, de-duplication errors, and oversplits due e.g. to spikes that are large on multiple channels (leading to variability in the assigned “primary channel” for these spikes), this first-pass estimate of templates typically requires some post-processing before we proceed with the deconvolution step. We outline two key post-processing steps below.

#### 2.8.1 Removing collision units

Despite the denoising and triaging steps described above, some units extracted in the first clustering pass remain dominated by collisions. That is, the unit template shape is a combination of 2 (or more) units. We can remove many of these units using one of two approaches.

First, recall that all spikes are aligned on their primary channel prior to the clustering stage. Therefore, if a template on its primary channel is significantly off-centered compared other templates, then it is considered as a collision unit and removed.

Second, we could look directly for “collision templates” — i.e., templates that can be well-modeled as a weighted superposition of two or more other templates. However, we found that this strategy was ineffective, since small timing variability between the two colliding spikes can quickly lead to distortions in the shape of the mean of these collided spikes. Instead we found that it was more productive to examine the variability of the spikes in each unit, rather than just the mean: small shifts in the timing of one spike relative to the other in a collision can lead to significant differences between waveforms, leading to large variability that we can detect reliably. We model spikes as the superposition of a template and a background noise term; thus, the covariance structure of a temporally isolated^§^ non-collided spike around its template is assumed to be the same as the covariance of the background noise. In addition, small misalignments of the spike (at resolution below the voltage sampling frequency) can also increase the variance. To address the issue, the maximal additional variance due to misalignment is estimated and incorporated in the variance bounds. This extra variance due to misalignment is estimated numerically by uniformly jittering the template with (−1,1) time step shift. Thus, we can search for units with higher-than-expected variability; we find that many of these units are clear collision units and can be discarded. See Figure 6 for an illustration.

**Figure 6:**
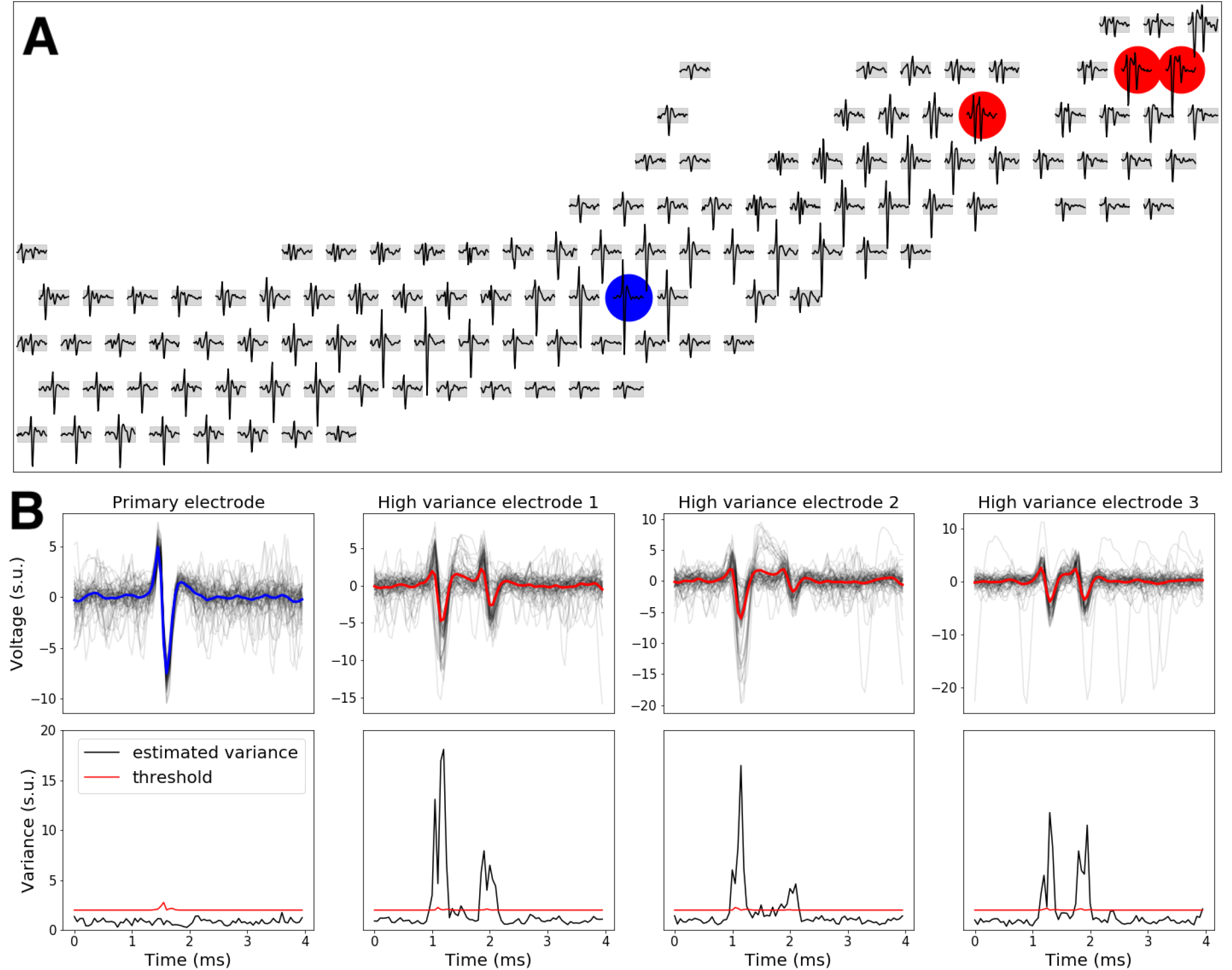
An example collision unit. (**A**) The template of the collision unit. Blue dot indicates the primary electrode (with the largest template), and red dots indicate the top 3 electrodes with the highest variability. (**B**) Top: raw spikes (gray traces) and templates (color) at the primary electrode and the three most-variable electrodes. Bottom: the estimated variance of spikes (black) and the threshold (red) used to detect overly-variable units. Due to small differences in the timing of one collided spike relative to the other, the variability of collided waveforms tends to be significantly elevated.

#### 2.8.2 Template de-duplication

We find pairs of duplicate templates by computing the maximum absolute difference (over time points and channels) between the aligned templates. Both absolute and relative differences are used to measure the closeness of two templates; if the maximum absolute difference is less than about 1 (absolute scale) or 10% of the size of the bigger template, we conclude that the templates match. (For low-firing-rate units, we compute templates using denoised spikes, since with few spikes the raw template may be corrupted by outliers due to collisions.)

Once a list of pairs of duplicate units are determined, the unit with more spikes is kept and the other is discarded, since the unit with more spikes will typically have a cleaner template.

### 2.9 Deconvolution

Once stable neuron templates are obtained, we run a deconvolution step on the raw voltage data to infer the spike times corresponding to these templates. This step is able to recover spikes that were triaged (due to collisions) or killed in the de-duplication step described above. A number of deconvolution approaches have been described in the literature (Franke et al., Carlson et al., 2013, Pillow et al., Ekanadham et al.). Our starting point is the matching pursuit approach used in (Pachitariu et al., 2016); this is a greedy method that essentially subtracts spike shapes from the raw data one at a time until a convergence criterion is reached. We improve this method in three key directions described below.

#### 2.9.1 Low-rank-plus-shift model for templates

Pachitariu et al. (2016) use a simple low rank model for templates (where the template is represented as a channelby-time matrix). This leads to faster computations (since the deconvolution time is dominated by convolution calls, and the number of required convolutions can be reduced by exploiting this low-rank template model) and can also reduce some noise in the templates, if the templates can be well-approximated by a low rank matrix.

Unfortunately, we found that this simple low-rank approximation is highly inaccurate for units with long axons, for which the spike at the soma can be significantly shifted in time compared to the spikes observed on axonal channels. In addition, as discussed above, many raw waveforms are corrupted by collisions, and therefore simple averaging of these raw waveforms can lead to noisy templates, leading downstream to noisier residuals in the matching pursuit algorithm and worse deconvolution results overall.

Instead we found that performing an alignment step (temporally aligning the templates on each channel) before computing the low-rank factorization led to significantly improved results (Figure 7). The goal of the alignment is to increase co-linearity of template matrix rows, therefore reducing SVD residual errors. To this end, for each neuron we minimize ℓ_2_ distances between the template on each channel or its negative (whichever leads to a smaller distance) and the primary channel template. In addition, we clean the waveforms (see section 2.10.1 below) before averaging to estimate templates, and then force the absolute value of the template to smoothly decrease monotonically to zero outside of a 2 millisecond window; both of these steps help reduce outliers prior to the align then SVD steps. Finally, we compute templates using isolated spikes (with interspike intervals greater than the width of the template), to avoid edge artifacts due to contributions from burst spikes.

**Figure 7:**
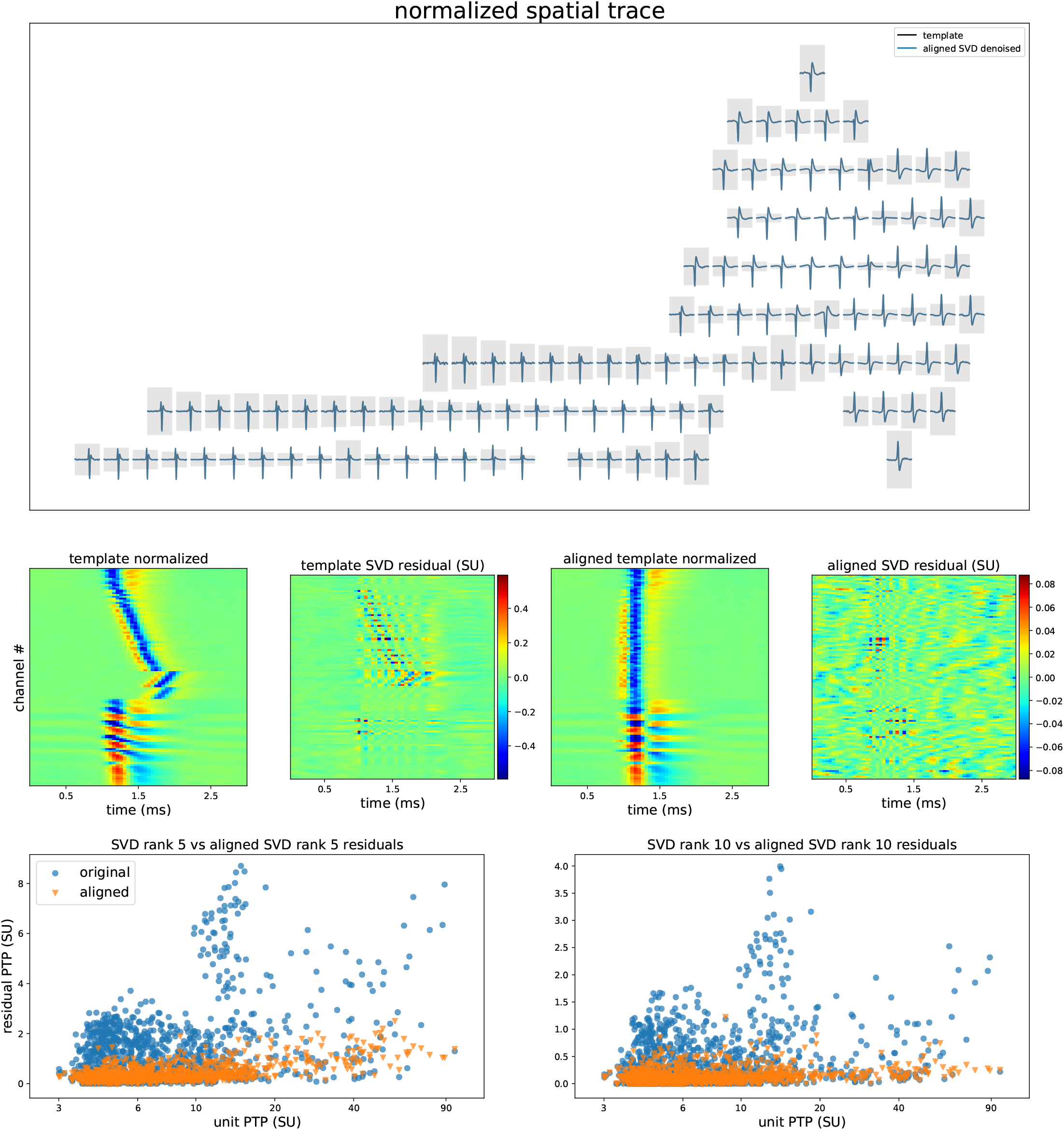
Temporally aligned low-rank template model. (**Top**) An example unit, showing that the raw template is well-summarized by its low-rank SVD reconstruction following temporal alignment, i.e., the blue and black traces show a high degree of overlap. Template plot conventions as in Figure 6. (**Middle row**) Comparing the SVD residual with and without alignment. First panel shows the raw template, normalized per channel; note that the peak time varies by about a millisecond across the different channels. Second panel shows that a low-rank SVD approximation leaves behind significant residual structure. Third panel shows aligned normalized template, and fourth panel shows that aligned SVD residual is significantly smaller than unaligned. (**Bottom**) Summary of residual errors across all templates in one dataset. For both rank 5 and 10 SVD approximations, aligning leads to significantly reduced residual error.

#### 2.9.2 Spline interpolation within deconvolution

In matching pursuit, first we identify putative spike times from a given cell and then we subtract away the corresponding template from the raw voltage trace. For large spikes, small timing differences between the spike and the corresponding template can lead to large residual errors if we naively subtract the template away; significant residual errors can accumulate even if these timing differences are below the acquisition sampling rate (20 KHz for the retinal data analyzed here).

This subsample-resolution subtraction issue has been previously addressed, e.g., in (Ekanadham et al.), who introduced an iteratively-reweighted L1 minimization approach. We implemented a simpler, more computationally efficient B-spline method for finding the optimal spike times at subsample resolution, then interpolating the template accordingly, and subtracting out this interpolated template. These steps are performed on GPU, leading to negligible speed differences compared to naive subtraction.

Figure 8 shows examples of the post-deconvolution residuals from large amplitude spikes. The interpolation approach leads to a substantially lower residual compared to naive greedy subtraction, and can thus yield substantially lower residuals, reducing deconvolution errors in the following iterations.

**Figure 8:**
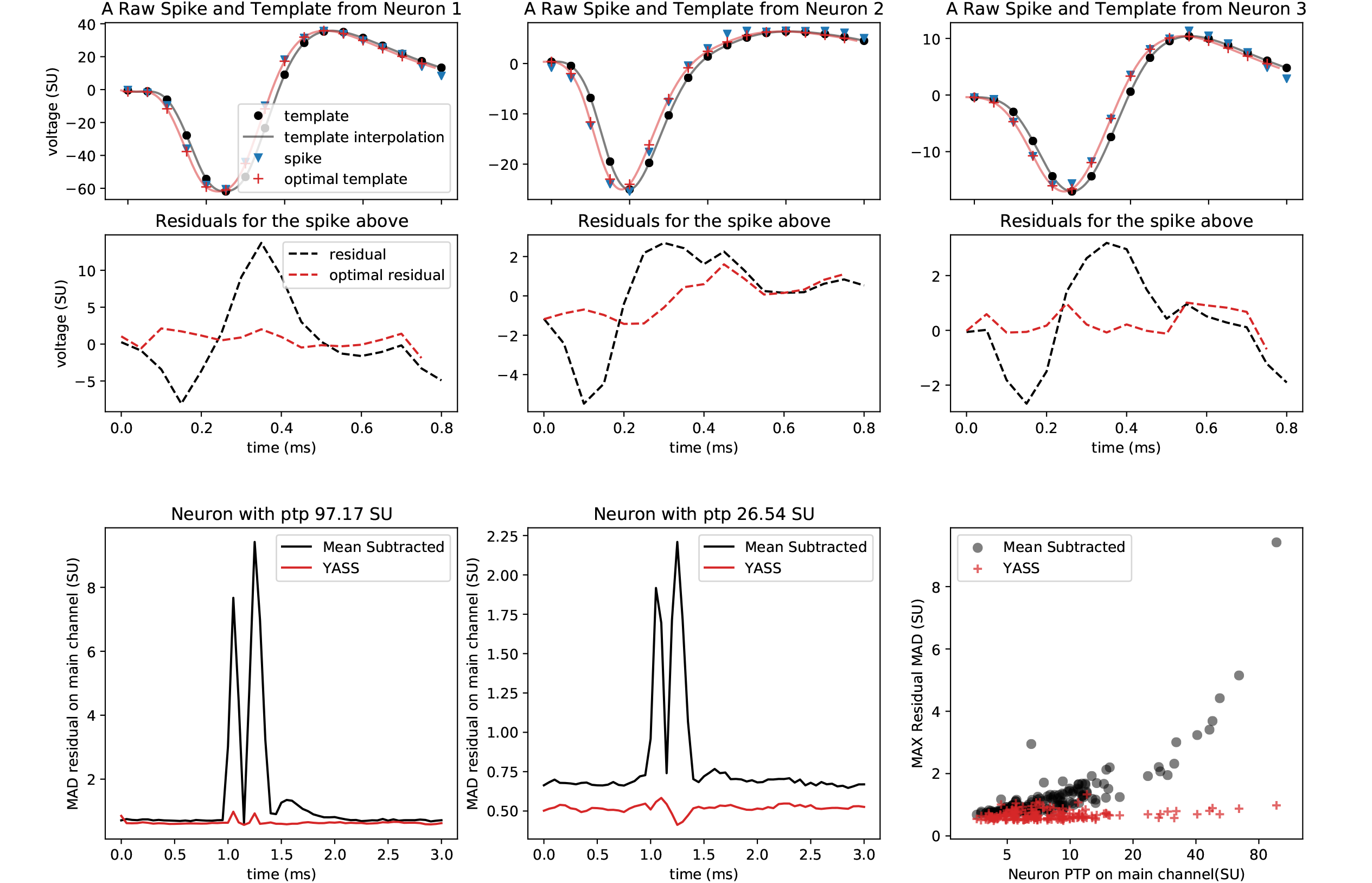
Super-resolution interpolation and subtraction improves deconvolution. (**Top row**) Three raw spikes from three distinct neurons are shown on the main channel of their respective neurons alongside neuron templates, their interpolations, and optimally time shifted and sub-sampled templates to achieve the smallest residual. Discrete signals are represented by scatter plot while interpolations are solid lines. (**Second row**) Residuals are shown when templates and optimal templates are subtracted from the raw spikes above. In all these cases the raw residuals have significant amplitude and can be mistaken as spikes from other neurons, while the optimally-aligned residuals are significantly smaller. (**Third row left & middle**) MAD of post-deconvolution residuals (in standardized voltage units) for spikes of two large neurons (“ptp” abbreviates the maximal peak-to-peak height of the spike template). Naive subtraction leaves significant residual variability behind (black traces); conversely, residuals are substantially smaller when super-resolution alignment and interpolation of the template prior are applied prior to subtraction (red trace). (**Third row right**) Residual variability using naive subtraction (black) scales with unit PTP, but with interpolation the residual error stays uniformly low (red; each symbol corresponds to a single cell).

#### 2.9.3 Reducing greedy deconvolution error through Iterative Coordinate Descent (ICD)

Matching pursuit is a greedy method: it subtracts spikes away one at a time, and never “reverses course”: e.g., it might become visually clear after subtracting more spikes away that an early spike was subtracted mistakenly, but simple greedy matching pursuit is unable to correct these errors. Thus it is natural to incorporate “reversal” steps into the algorithm — i.e., instead of always subtracting spikes away, we could also propose adding previously-removed spikes back in, and perhaps subtracting different spike times away in a second pass through the data. This combination of subtraction and addition steps is a well-known approach in the sparse regression literature (Zhang, 2011).

To make this idea slightly more concrete, matching pursuit can be cast as a greedy optimization of a squared-error objective function, where we are optimizing over the space of binary matrices indicating the presence or absence of a spike from each cell in each time bin. One simple way to improve on matching pursuit is to replace this greedy optimization with iterative coordinate descent (ICD) on the elements of this binary spiking matrix, using the same squared-error objective function. To implement ICD here we simply augment matching pursuit with steps for changing spike times or adding spikes back in if they have been subtracted away previously. These steps can be easily incorporated into the existing matching pursuit GPU code; in practice we just end up taking an additional couple passes through the data in the deconvolution step.

While ICD does not correct all false negative errors, it decreases them in almost all cases relative to greedy deconvolution, in some case by an order of magnitude (Figure 9A). Empirically, we find that this ICD approach can increase true-positive rates by a few % for many units, while significantly decreasing false-negatives for many neurons.

**Figure 9:**
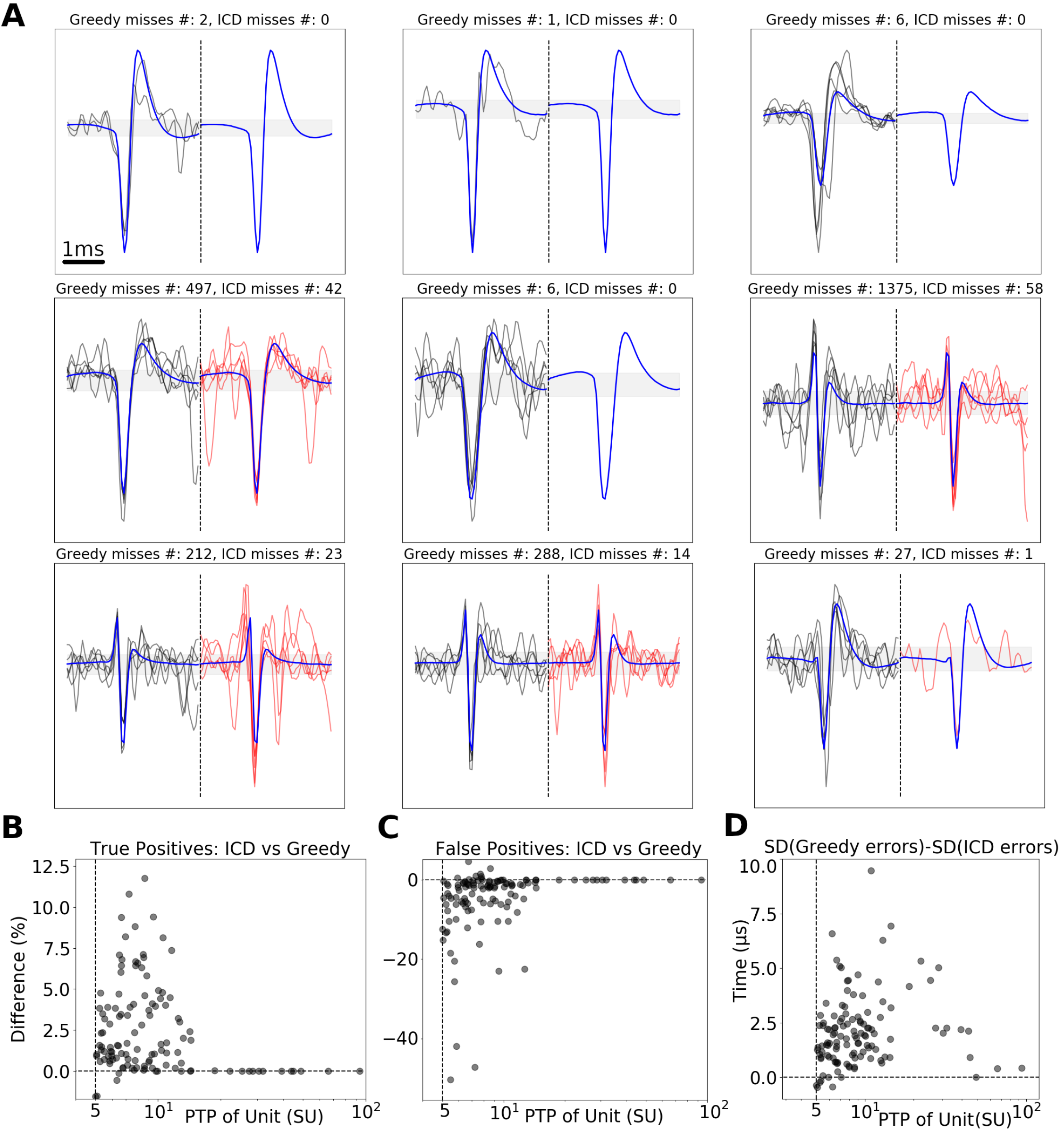
Comparing greedy vs iterative coordinate descent (ICD) deconvolution. **(A)**: Examples of spikes recovered by ICD but missed by greedy deconvolution (and vice versa). All comparisons here performed on synthetic data with known ground truth spike times (see section 2.13 for a description of synthetic data generation). Blue indicates the true underlying (noiseless) spike template; black is an example of an observed noisy voltage trace containing the spike that greedy deconvolution missed but ICD caught; red indicates traces for which ICD missed the spike but greedy deconvolution accurately captured the spike (up to a maximum of 5 traces are plotted). Note that the miss rate is often much lower for ICD. **(B-D)**: ICD improves true positive rates on the order of a few % **(B)**, decreases false positive rates by up to 50% **(C)**, and modestly improves the timing precision of detected spikes, as measured by the standard deviation (SD) of the inferred spike times **(D)**. In each case, differences between greedy versus ICD are most significant for smaller units.

### 2.10 Post-deconvolution merging and splitting

The deconvolution step can recover spikes that were dropped in the triage or de-duplication step. In addition, much of the “noise” we observe in each spike waveform is in fact due to collisions with other spikes; once we have determined these other spike times we can subtract out the corresponding spike shapes and therefore decrease the effective noise level. After this “spike waveform cleaning” step, in turn, some new clusters that were not visible before this “cleaning” step can become easily separable. Therefore we re-run the clustering step on the “cleaned” waveforms, re-splitting and merging as necessary to improve the overall cluster assignments. We describe each of these steps below.

#### 2.10.1 Post-deconvolution cleaned spike waveforms

As a first step we need to clearly define these cleaned waveforms, and then describe efficient methods for computing these objects. The deconvolution step returns spike times for each cell, along with the *residual* (i.e., the original raw voltages, minus the contribution of each spike that was subtracted away during deconvolution). (Ideally, if we are capturing all the spikes, this residual will resemble noise, in which no clearly discernible spikes remain.) The *i*-th “cleaned” spike waveform is defined as the *i*-th spike waveform minus the contribution of all the other spikes with indices *j* ≠ *i*. To compute this efficiently, we can simply rewrite this as the residual (which subtracts all of the spikes out, including i) plus the template for spike *i* added back in. Thus we can compute each cleaned waveform rapidly, in an embarrassingly parallel manner over all the spikes. See the top row of Figure 10 for an illustration.

**Figure 10:**
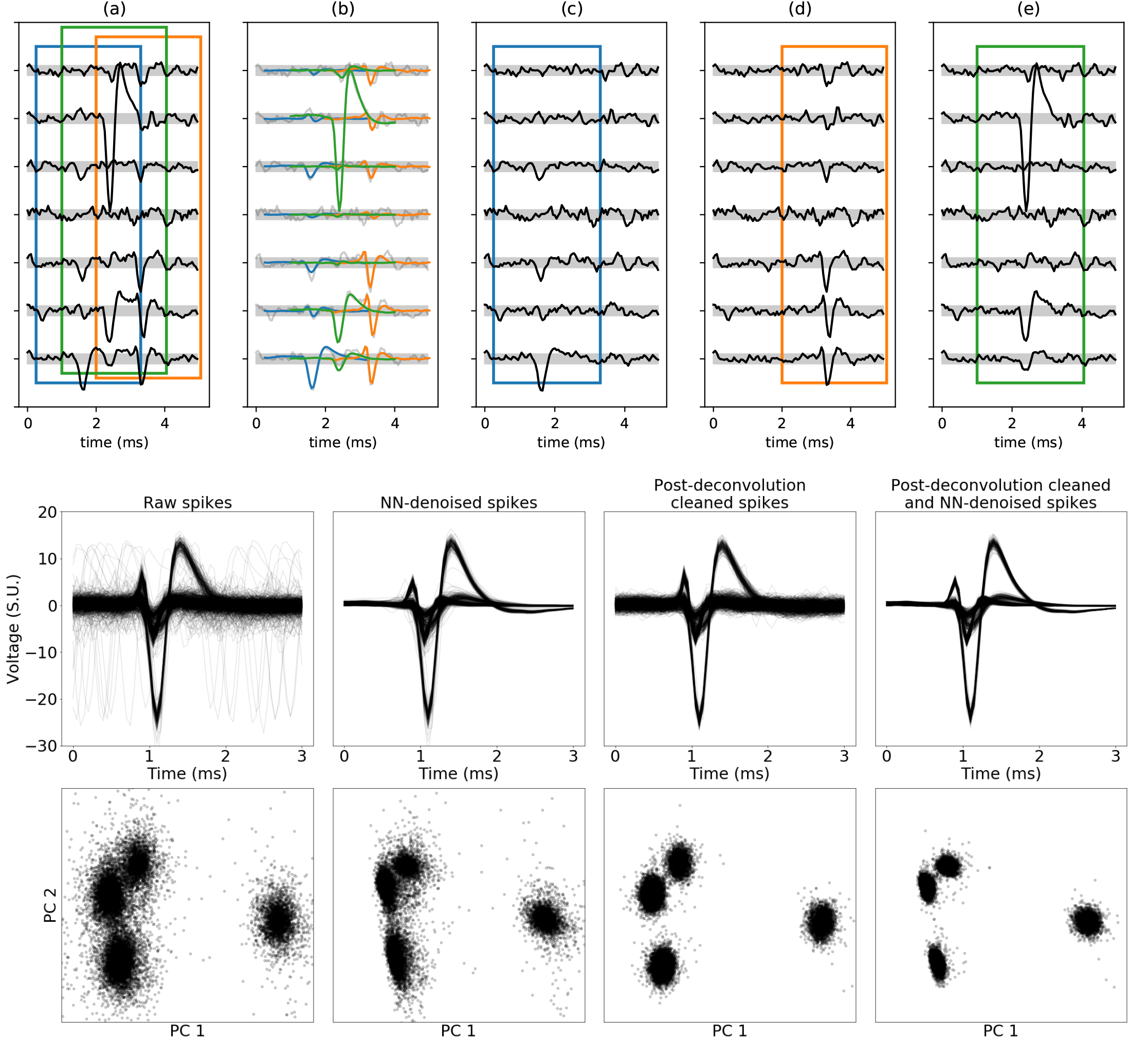
Post-deconvolution spike waveform cleaning. **(a)** A 5ms 7 channel raw voltage recording reveals the presence of multiple spikes. **(b)** Three spikes are identified by deconvolution and are depicted by inserting each template (colored traces) at the times identified by deconvolution. **(c, d, e)** “Clean spikes” for each of the three spikes are obtained by superimposing the template shape with the residual voltages after all the other identified neuron spikes are removed from the raw record. **Middle row**: Raw, NN-denoised, post-deconvolution cleaned spikes, and cleaned then NN-denoised spikes from a single channel. **Bottom row**: PCA projections corresponding to the waveforms from the middle row. Note that post-deconvolution spike cleaning and NN-denoising both significantly improve the effective SNR of the data.

#### 2.10.2 Post-deconvolution reclustering

We do not expect the original clustering step described above to split all the cells perfectly: due to the inherent noisiness of the raw data, the high collision rates, and the stochasticity of our clustering algorithm (which includes some randomized split-merge moves), some neuron templates (albeit usually a small minority) can be missed, incorrectly merged, or split by our pipeline.

After deconvolution and post-deconvolution cleaning, we have more spikes than what was used in the original clustering step (because now we have recovered spikes that may have been triaged or killed during the first pass), and the cleaned spikes show significantly less variability than the raw data (Figure 10, lower panels). Therefore, a re-clustering step run on this post-deconvolution data has the potential to correct some errors that may have been made in the first pass.

The deconvolution step has already grouped spikes according to the unit each waveform was assigned to. We run our clustering pipeline individually on each of these groups. Because most of these spike assignments from the deconvolution step are already fairly accurate, this reclustering step tends to be significantly faster than the original per-channel clustering run; in most cases this re-clustering step merely reconfirms the original template identity, with occasional splits identifying additional neurons. In addition, because most outliers are due to collisions, and the deconvolution step has already resolved most collisions, we found that it is unnecessary to perform triaging during this re-cluster step.

#### 2.10.3 Merging units using cleaned spikes

As in the first clustering pass, it is necessary to perform a final merge step following the re-cluster step to try to fix any oversplits that may have occurred. Here we use an approach based on linear discriminant analysis (LDA) applied to the cleaned waveforms.

We start by computing a candidate list of pairs of units to be considered for merging; any pair of units with aligned templates that are sufficiently close in ℓ_2_ distance are added to this list. For a candidate pair of templates, we then select a subset of spikes (usually up to 2,000), compute cleaned spikes, and whiten the resulting waveforms. To avoid overfitting, we compute a separable (Kronecker) approximation to the whitening matrix (as in (Pillow et al., Ekanadham et al.). Next, we project the whitened cleaned spikes onto the difference of the two templates (restricted to the union of the active channels of both units). (This projection is equivalent to LDA if the whitening matrix is a good approximation of the inverse square root of the covariance of both units.) If the marginal distribution of the resulting one-dimensional data is approximately unimodal (according to a Hartigan dip test (Hartigan et al., 1985)), we merge the units; see Chung et al. (2017) for a related approach but applied to raw voltage data. (We merge pairs greedily, merging pairs with smallest template distance first.) Figure 11 visualizes the procedure for two pairs of neurons and demonstrates why it is beneficial to use cleaned spikes as opposed to raw spikes in the analysis.

**Figure 11:**
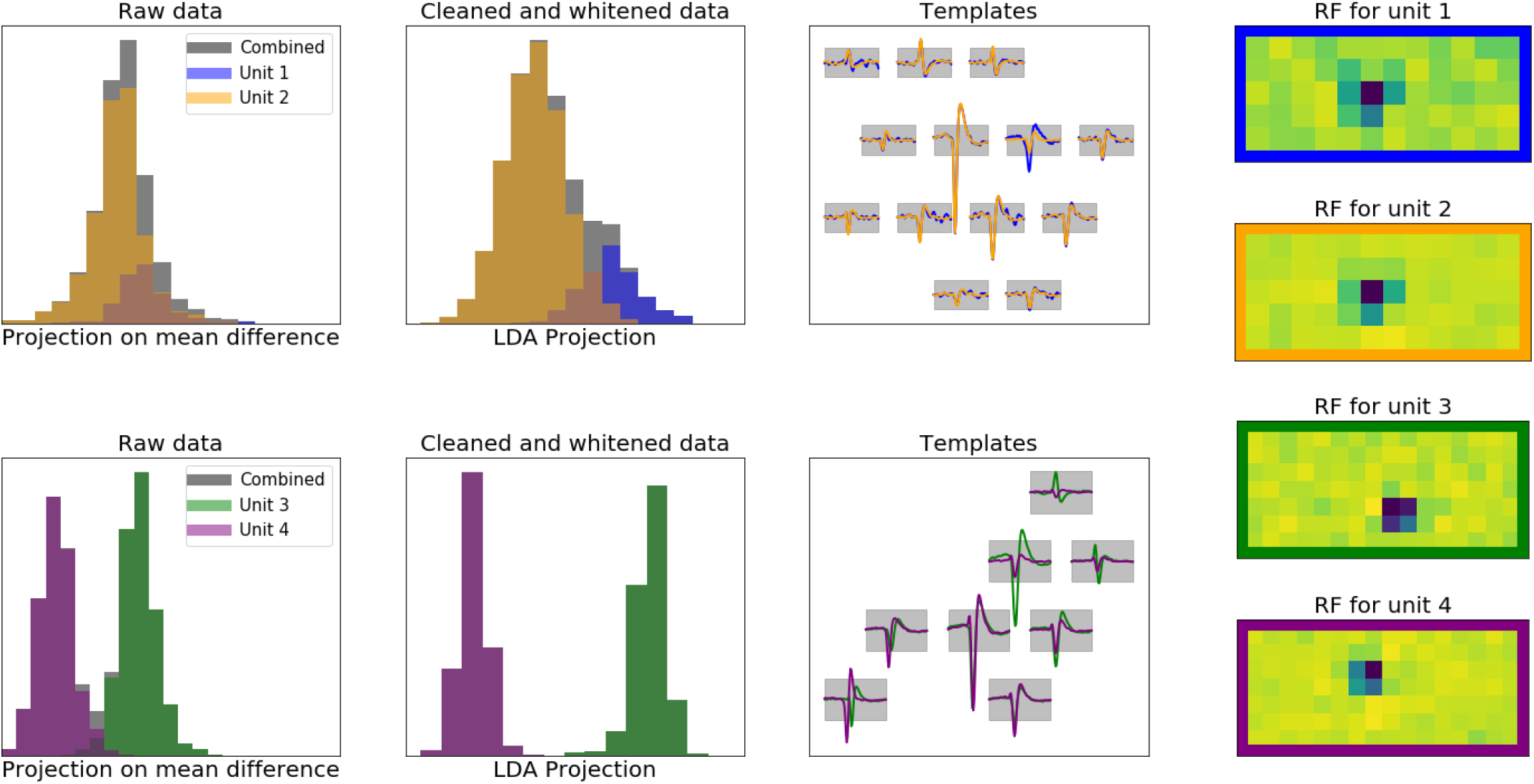
Post-deconvolution merge using cleaned spike waveforms. Each row illustrates the process for a different pair of units whose templates are close enough to be candidates for merging. **First column**: for each unit, we project the raw waveforms onto the difference between the two templates and plot the resulting histograms. **Second column**: same, but projecting the post-deconvolution cleaned waveforms onto the linear discriminant analysis (LDA) direction; note that this improves the separation. **Third column**: the templates of each pair of units. **Last column**: receptive fields (RFs) for all these neurons; note that these RFs are not used in the merge algorithm, but are shown here for verification purposes only. Using LDA applied to clean spikes, our algorithm suggests merging templates 1 and 2 while avoiding a merge of templates 3 and 4. This decision is supported by the corresponding RFs: RFs 1 and 2 are similar (though RF 1 is much noisier due to having fewer spikes than neuron 2), while RFs 3 and 4 are clearly distinct.

### 2.11 Drift handling

In the retinal recordings considered here we observe spike amplitude changes on the order of 1% (or less) of PTP height per minute (Figure 12). We correct for this drift by implementing a model with two stages: (1) we obtain neuron templates from the beginning of a recording (first 5 minutes); and (2) update the templates periodically after every new batch of deconvolved data (in batches of size 5 minutes). (See Pachitariu (2019) for a related method.)

**Figure 12:**
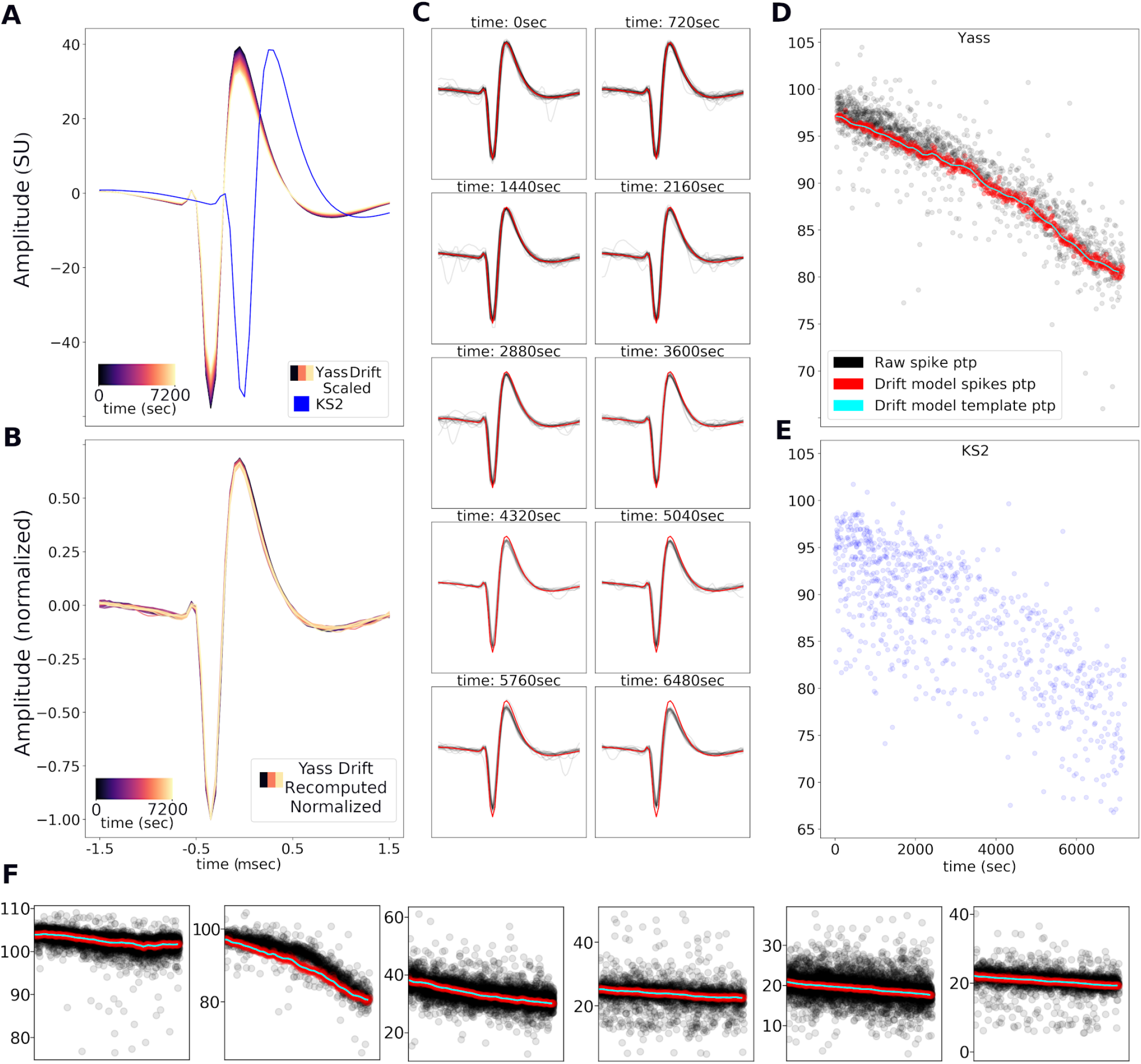
Drift tracking via template scaling. **A**. Drift model example: a template is scaled to match the drifting size of a neuron observed over a period of 2 hrs (magma colors; matching template from Kilosort is in blue and was computed using 5 minutes of data). Note the YASS template amplitude decreases by 15%-20% over the duration of the 2 hr (7200 sec) recording. **B**. Testing the stability of the template and rank 1 approximation: templates recomputed (and normalized) every 2 minutes for 2 hrs using deconvolved spikes (i.e. each template represents the mean of the spikes deconvolved in a 2 minute period). Note that the normalized template is nearly identical over time, suggesting scaling is an adequate model of drift. **C**. Template from the first 5 mins of recording (red) with deconvolved spikes (10) across 2hrs of recording. The change in individual spike size (black traces) compared with the starting template is observable at the single spike level. **D**. Drift model tracks changes: PTP of raw deconvolved spikes (black), PTP of spikes selected for the drift update (red) and PTP of the drift model (cyan). This shows that the drift model is tracking drift over time despite periodic updates (e.g. every 2 mins). **E**. PTP of spikes deconvolved for the same neurons by Kilosort. **F**. Additional examples of drift tracking; conventions as in panel D, but applied to different units.

To update the templates we exploit the low-rank model described in section 2.9.1: i.e., instead of updating the raw templates (which are specified by many parameters, leading to potential overfitting in units with low spike counts) we update the parameters of the low-rank-plus-shift model directly, using exponential weighted updates. Furthermore, in these recordings we found that templates change largely in amplitude rather than shape (See Fig 2.11, A-C). Accordingly, in each channel we compute the updates by scaling the starting templates based on the weighted average of the spikes in the previous deconvolved batch and the following batch. Due to the high collision rate present in raw spikes, we base the update solely on cleaned spikes falling within +/-20% of the starting template (see red scatter plot in Fig 2.11 D,E). While this limits the rate of change in the drift model, we found that it accurately tracks the change observed in our MEA recordings.

To add templates of intermittently-active neurons that might not have been active in the the initial batches of data, a simple split step is performed in each new batch. For each unit, the PTPs of cleaned spikes are computed in each channel and we test the uni-modality (Hartigan et al., 1985) of a one-dimensional PCA projection of these PTP values. If the diptest p-value is below a threshold, the unit is split using a two-cluster EM algorithm. The same method is applied recursively to ensure the uni-modality of all clusters. To reduce oversplits, the split units are accepted only if their firing rate is bigger than 0.5Hz or if the ratio of its firing rate compared to the original unit is bigger than 0.15. Any split units that are classified as likely duplicates (section 2.8.2) or with low PTP units (less than 2.5 standardized units) are also removed.

After completing a forward pass through all batches (adding in new templates as needed), we perform a backwards deconvolution (to properly assign spikes to units that might have only been identified in the middle of the recording), updating the templates as we pass through batches in reverse order, but not performing any additional split steps. Then we drop any units that have low firing rates (less than 0.2Hz), small PTP (smaller than 2.5 standardized units), or low average soft assignment score (see section 2.12). We then run a final forward deconvolution to assign spikes to the remaining units, with no further template updates or split steps.

### 2.12 Soft assignment

For lower-SNR units, there will invariably be some borderline spikes for which the correct assignment is unclear. In addition, even high-SNR units may contain some outliers (e.g., due to deconvolution or clustering errors). Therefore, as a final postprocessing step we use the cleaned waveforms to compute three soft assignment scores:

1. Soft noise assignment: for the smallest units (according to the ℓ_2_ norm of the template, restricted to the unit’s active channels), we run the detection neural network on the cleaned waveforms and record the output scores; these can be used downstream by the user to perform a soft-classification into noise versus signal waveforms.
2. Outlier scoring: for all units, we compute the ℓ_2_ norm of the whitened cleaned waveforms (using the approximate Kronecker whitener described in section 2.10.3); this score can be used to soft-classify outliers for each unit.
3. Soft pairwise assignment: for the pairs of units that are closest (according to the ℓ_2_ norm of the difference of the aligned templates, restricted to the union of the two units’ active channels), we compute the LDA projection (again as in section 2.10.3), estimate the density of each unit in the resulting one-dimensional projected space, and compute the likelihood ratio. For each pair, if one unit has an average soft assignment score below a threshold, we consider the unit a duplicate and remove it.

### 2.13 Synthetic and semi-synthetic data

For testing purposes it is invaluable to run spike sorting pipelines on both fully-simulated and “hybrid” simulated+real datasets in which ground truth is either fully or partially available. For rapid testing purposes, we generated simulated datasets on 49 channels. We estimated templates from a held-out dataset, and then applied randomly sampled scale factors to these templates. For fully-simulated datasets, we selected about 150 templates and generated spatiotemporally correlated noise as the background signal (this noise was in turn generated by convolving many independent white noise traces with randomly chosen templates, then scaling and averaging the resulting spatiotemporal noise together). For “hybrid” datasets, following (Rossant et al., 2015, Pachitariu et al., 2016), we chose about 20 templates and added spikes with these template shapes to a separate real dataset which served as the “background” signal. Next we randomly sampled firing rates uniformly on (1, 30) Hz, sampled Poisson spike trains with these rates, and added the selected templates (with sub-sample jitter and interpolation) at the selected times.

## 3 Results

### 3.1 Basic characterization of YASS output

Figure 13 provides some basic statistics characterizing the output of YASS applied to three retinal datasets (number of units extracted, firing rate, etc.). As emphasized in Figure 1, due to the large number of spikes visible on each electrode, the collision rate is high in these datasets: for about half of the recovered units, about 20% of all spikes have a collision with a spike of peak-to-peak size ≥ 4 within 0.5ms.

**Figure 13:**
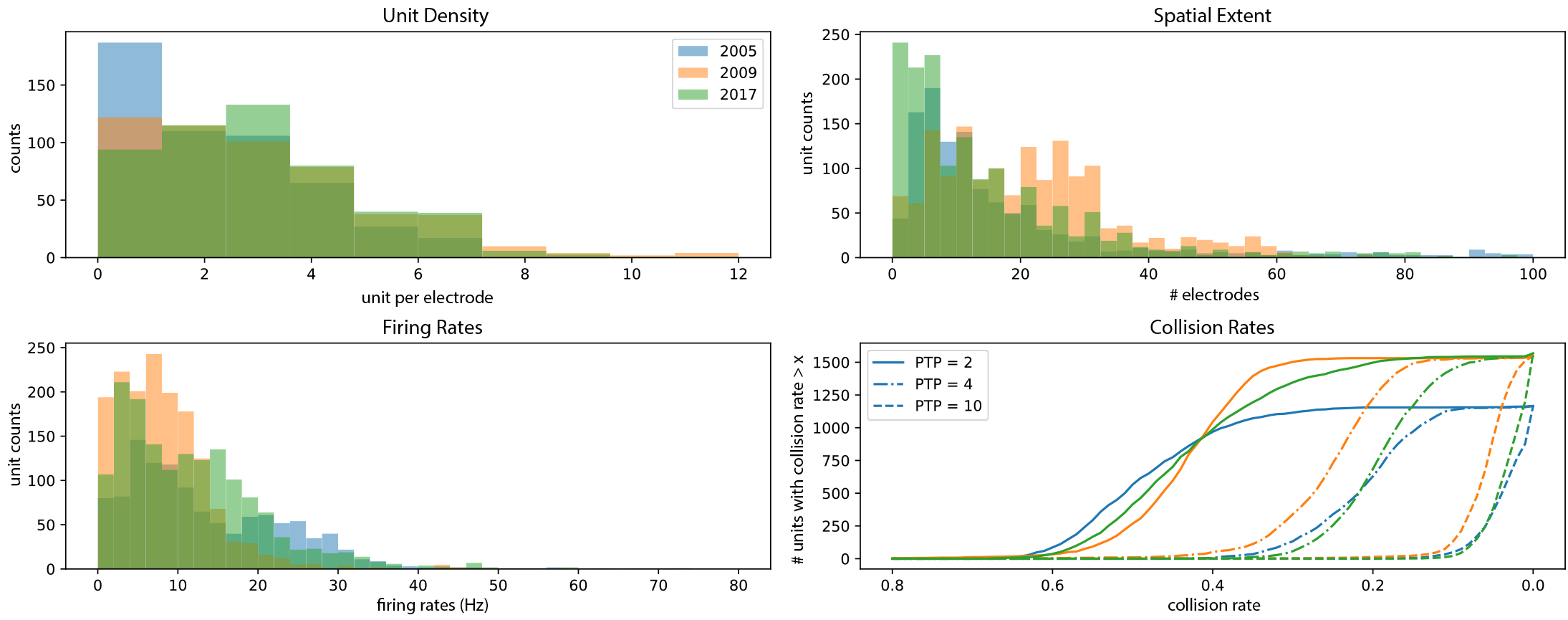
Basic statistics of units extracted from three retinal datasets. **First panel**: number of units extracted per electrode for three datasets. **Second panel**: Spatial extent of each neuron, measured by counting the number of electrodes on which the template PTP was greater than 2 (note, 2009 dataset contained a higher incidence of axons, leading to more units active on a large number of electrodes). **Third panel**: Estimated firing rates of all sorted neurons. **Last panel**: The rates at which spikes collide with other distinct signals of PTP size 2, 4, and 10 within a 0.5ms time window for the three datasets. Concretely, for about half of the recovered units, about half of all spikes have a collision with a spike of PTP size ≥ 2 within 0.5ms; similarly, for about half of the recovered units, about one fifth of all spikes have a collision with a spike of PTP size ≥ 4 within 0.5ms.

Figure 14 and 15 summarize several diagnostics that we found useful while iteratively improving the algorithms described here. Figure 14 illustrates one of our basic diagnostic tools: examination of multiple templates and receptive fields (RFs), divided by cell type. (Cell types were clustered using standard approaches, as described e.g. in Field et al. (2007).) Figure 15 shows further diagnostic plots used to examine the health of a single sorted unit, including cross-correlations and RFs of neighboring cells (where “neighbors” here can be defined in terms of distance between RFs, or templates, or in terms of the height of the peak cross-correlation between pairs of units); these are useful in determining whether a unit has been over-split or over-merged.

**Figure 14:**
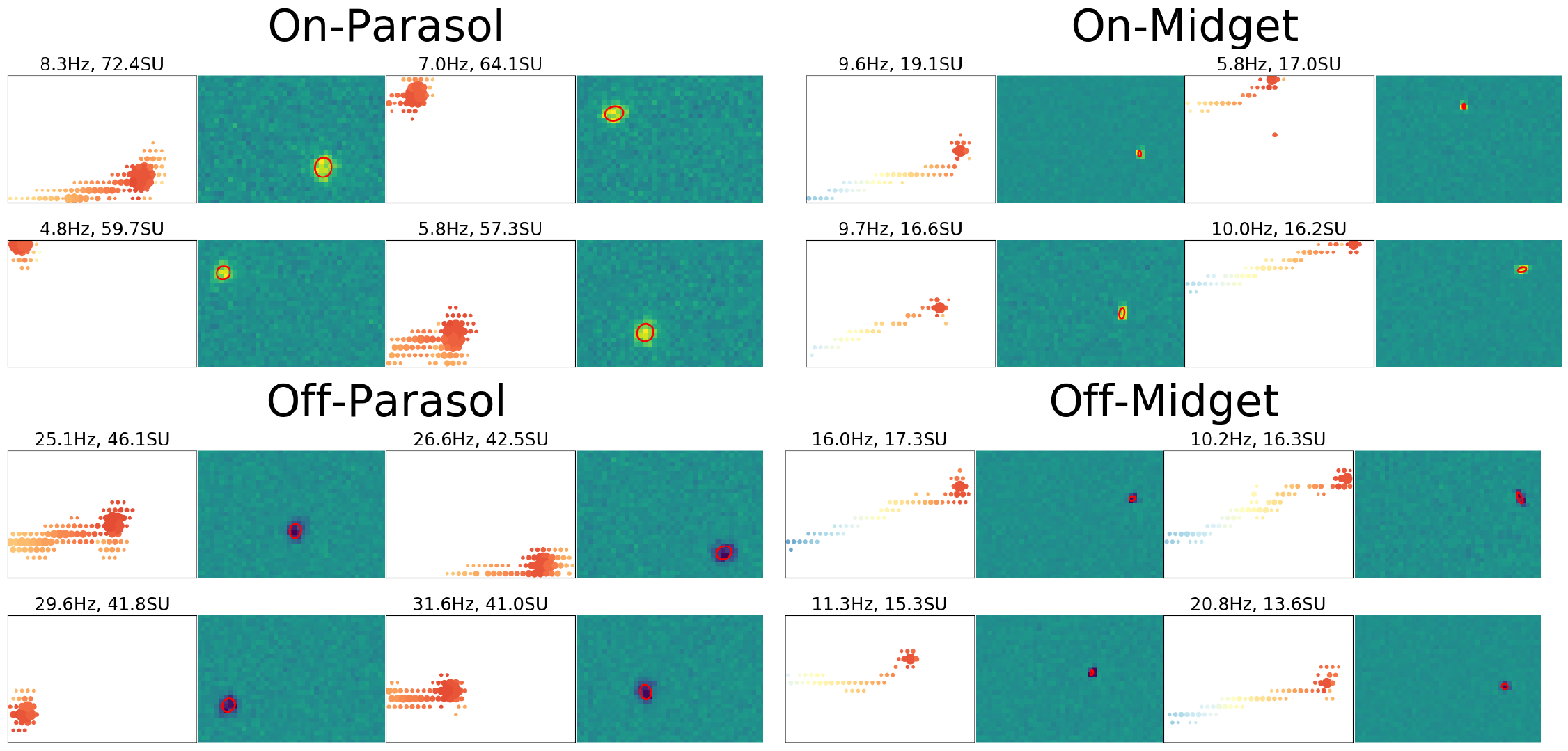
Spatio-temporal templates and receptive fields (RFs) from four major cell classes. **Left panels**: Spatial (filled circles) and temporal (colors) representation of templates for neurons sorted using YASS. Each dot represents the PTP of the waveform on the indicated channel (only channels with PTP > 2.0 are shown); warmer colors (orange) indicate earlier time (i.e. closer to spike time) and cooler colors (e.g. blue) represent channels active at later times. **Right panels**: Spatial RF of neurons corresponding to the templates shown in the left panels. Red contour shows Gaussian fits (c.f. Figure 16 below). Cell types were determined from RFs by standard clustering procedures (Field et al., 2007), with borderline cases determined by hand using a simple custom GUI.

**Figure 15:**
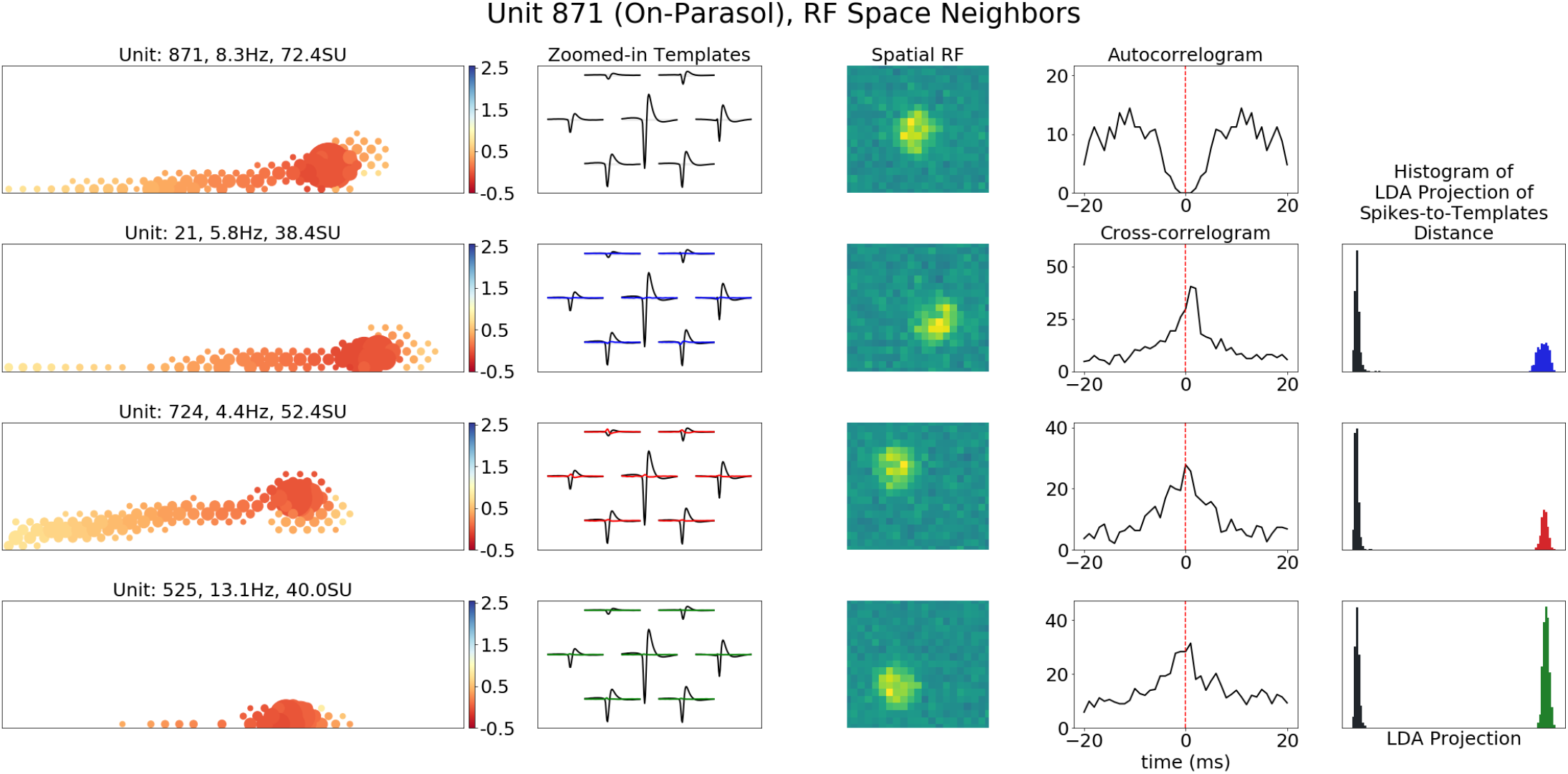
Example diagnostic plots for a single recovered unit. To examine the health of the extracted units we examine templates, RFs, and autocorrelations, along with corresponding plots for “nearby” units (where unit “neighbors” can be defined according to distance in RF space, template space, or other parameters such as the degree of correlation strength). Here we show an example of some of these analysis plots for an ON-parasol cell and its four nearest neighbors as defined by RF cosine distance. From left to right, the diagnostic plots show: template propagation profile across the 512 channel array (time since spike on primary channel in ms represented by color; firing rates and PTP in standardized units are provided in the panel titles); overlapping template plot of the main neuron on its primary and neighboring channels (black) and its neighboring cells (color); the (cropped) spatial RF; autocorrelogram and cross-correlograms with the neighboring units (in Hz); and histograms of spikes-to-template distance for all spikes in each neuron against each template after the LDA projection described in section 2.10 (more distance between the histograms indicates that the cells are better separated).

### 3.2 Comparisons to manual sorts and other automated spike-sorters

We have also evaluated spike sorting results using several other sorters: Kilosort (Pachitariu et al., 2016, Pachitariu, 2019), Spyking Circus Yger et al. (2018), Mountainsort Chung et al. (2017), JRClust and Ironclust (Jun et al., 2017b), and Herding Spikes Muthmann et al. (2015). Of this group, in our hands Kilosort consistently led to the best results on the retinal datasets considered here. Therefore in this section we will restrict attention to Kilosort.

Figure 16 is perhaps our main result: we show that on real MEA recordings YASS is able to recover much more of the underlying mosaic structure in the RFs of the four main retinal ganglion cell types (Field et al., 2007), compared to the output of both a highly laborious manual sort and the best current semi-automated method (Kilosort). These differences are particularly striking for midget cells, which tend to have smaller spikes compared to parasol cells, thus making their spikes more challenging to sort.

**Figure 16:**
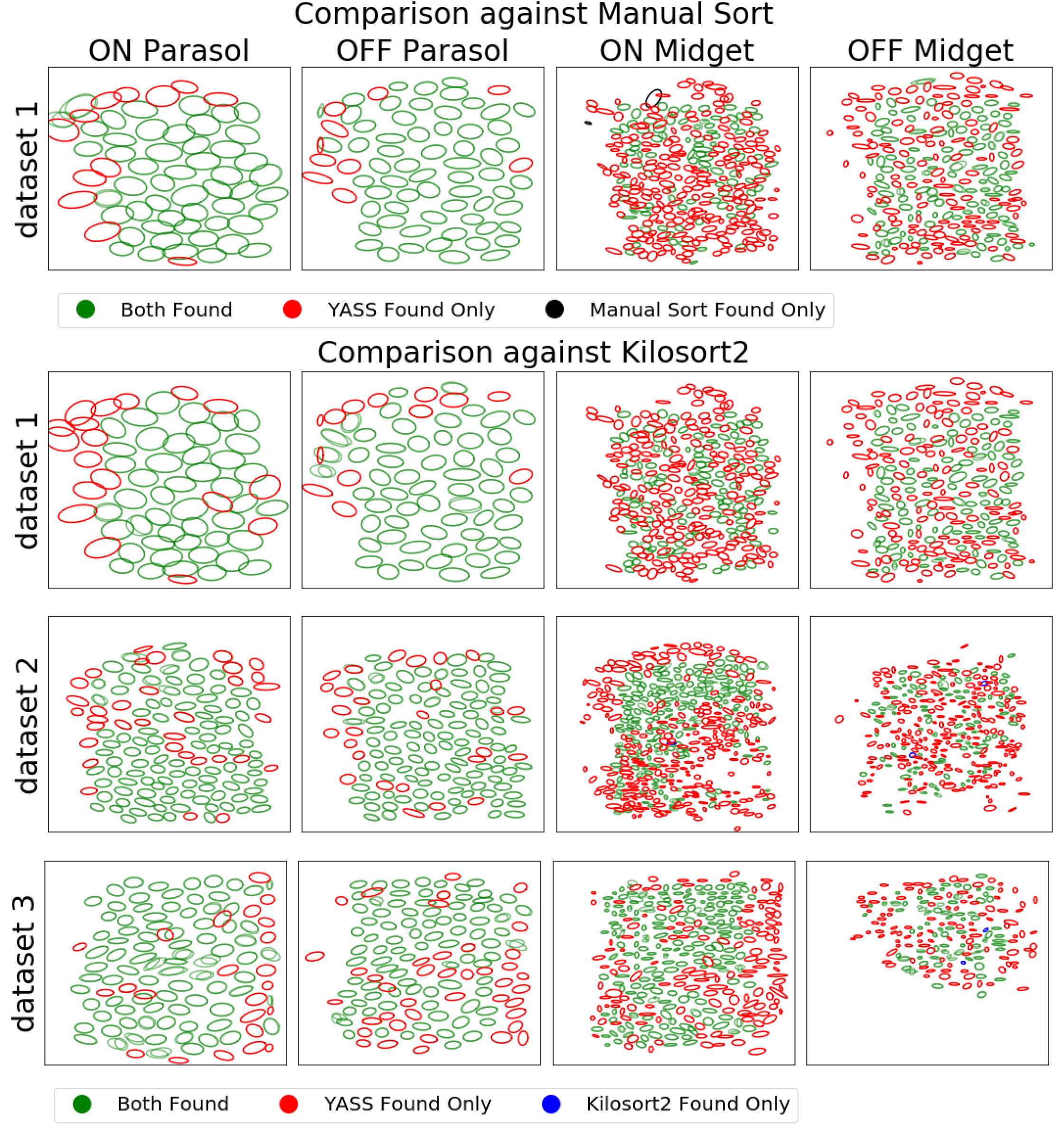
Receptive field contour analysis. Each ellipse indicates a Gaussian fit to the receptive field of an individual unit. (Top) RF contours of units extracted by YASS and a hand sorted result on one 512-electrode dataset. Green indicates units that were found by both YASS and the hand sort; red indicates units that were absent in the hand sort; black indicates units that were found in the hand sort but not by YASS. (Bottom) Contours of Yass and Kilosort2 on three separate 512-electrode datasets. YASS consistently recovers more cells with RFs, particularly midget cells.

Figure 17 illustrates a useful method for comparing between two spike sorters on real data, where no ground truth is available. For a given unit output by spikesorter A (e.g., YASS) we find the matching unit (or multiple units) output by spikesorter B (e.g., Kilosort). Then we run a spike-by-spike comparison to see which spikes were assigned to which unit; this comparison can help indicate when one spikesorter is mis-assigning spikes, or is over-splitting units, or is missing units. The caption of Figure 17 provides further details. We summarize the results of this analysis in Figure 18: we find that the extra spikes found by YASS are typically similar to the “consensus” spikes found by both sorters; conversely, the extra spikes found by the non-YASS spike sorter (either manual sorting or Kilosort) often look significantly different from the “consensus” spikes (as measured by the cosine similarity of the average spike shape in each group). For many units, YASS recovers tens of spikes per second that the other spike sorters missed.

**Figure 17:**
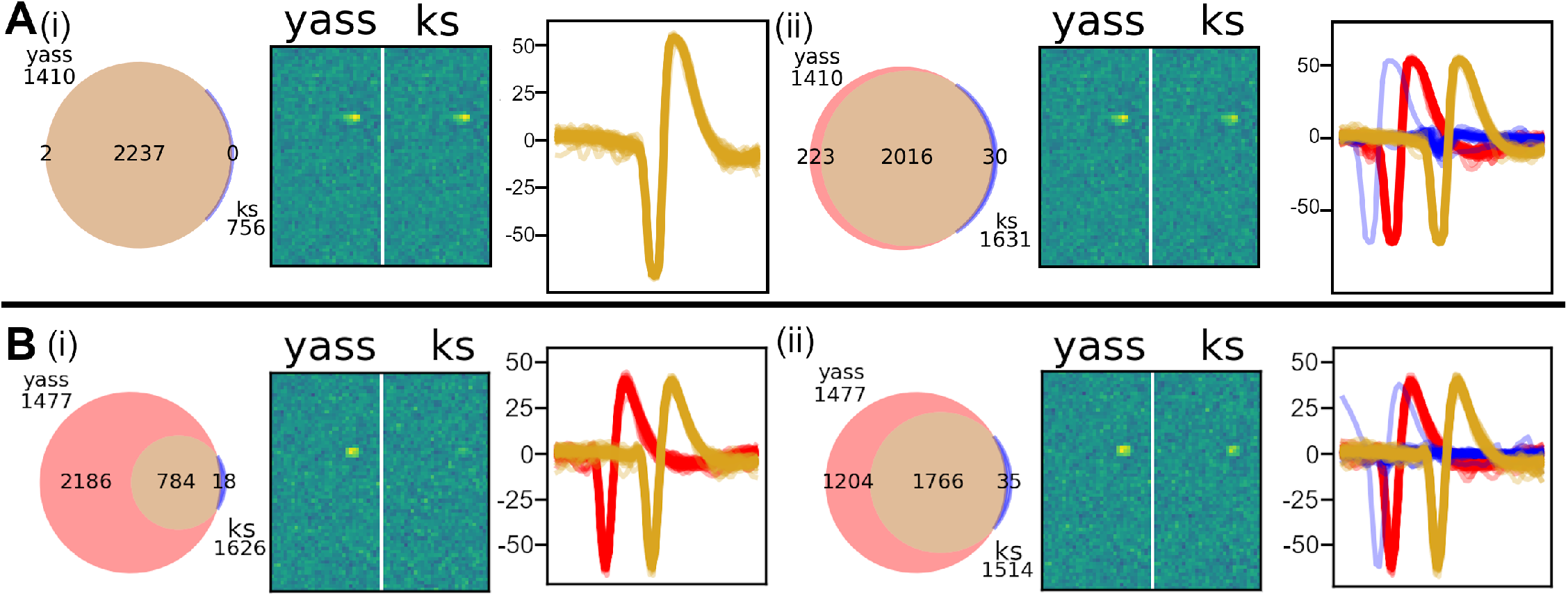
Pairwise per-unit comparisons of spike sorters. **A**. Example of an apparent Kilosort de-duplication error. **(i)**: (*Left*) Venn diagram for YASS unit 1410 (red) and Kilosort unit 756 (blue), showing an overlap of 2237 spikes (beige), with YASS identifying 2 extra spikes. (*Middle*) RFs for the 2 units. (*Right*) Spikes randomly selected from the intersection set. **(ii)**: (*Left*) Kilosort unit 1631 also matches YASS unit 1410, indicating that Kilosort unit 1631 is a duplicate of Kilosort unit 756. (*Middle*) RFs for the units (note the slightly weaker Kilosort RF, likely due to slightly fewer spikes). (*Right*) Spikes taken from the intersection (beige) look similar to additional spikes from YASS (red) but match poorly with many additional spikes from Kilosort (blue), indicating that Kilosort unit 1631 contains some incorrectly assigned spikes. Note that spikes from different groups are displaced horizontally for better visibility here. **B**. Example of a likely Kilosort oversplit error. **(i)**: (*Left*) Venn diagram for YASS unit 1477 (red) and Kilosort unit 1626 (blue) have an overlap of 784 spikes (beige), with YASS identifying 2186 spikes and Kilosort identifying 18 extra spikes. (*Middle*) RFs for the 2 units (note that Kilosort RF is relatively weak here). (*Right*) Spikes randomly selected from the intersection set (beige) and YASS (red) indicate that additional spikes identified by YASS are consistent with the spikes recovered in the (beige) intersection set. **(ii)**: (*Left*) Kilosort unit 1766 also matches YASS unit 1477. (*Middle*) RFs for the units (note the slightly weaker Kilosort RF). (*Right*) Spikes taken from the intersection (beige) look similar to additional spikes from YASS (red) but not most additional spikes from Kilosort (blue), again indicating that Kilosort unit 1631 contains incorrectly assigned spikes.

**Figure 18:**
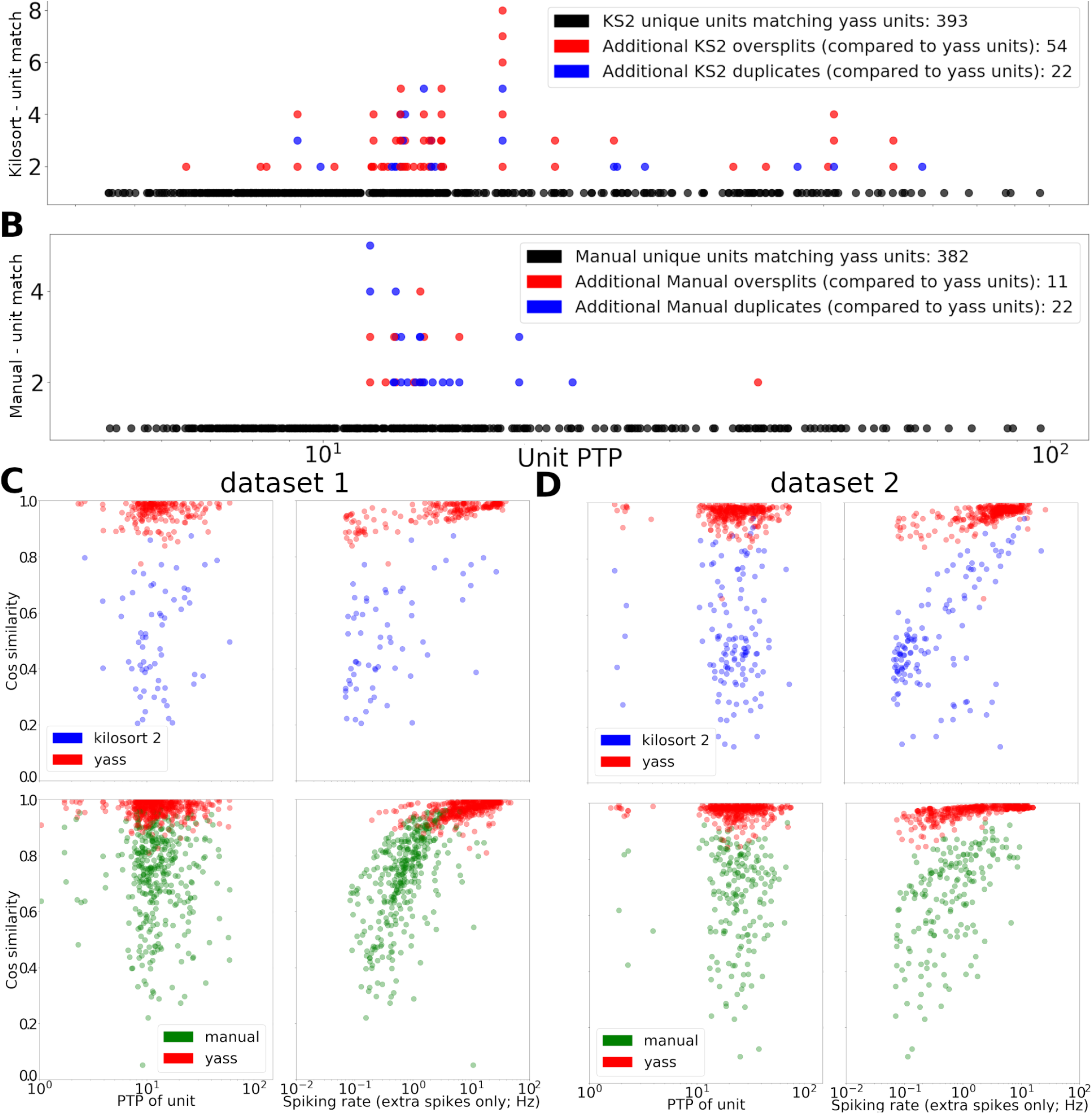
Summary analysis of oversplits, duplicates, and consensus across spike sorters. **A**. Summary of Kilosort units matched to YASS units. Most KS units matched well with YASS units (black dots) but there were many examples of oversplits and duplicates (as described in Figure 17; duplicates were defined as KS units that matched a YASS unit after another KS unit was matched; oversplits were defined as duplicates with fewer than two thirds of the YASS unit spikes). The y-axis indicates how many KS units were matched to a single YASS unit (see the legend for details). **B**: Same analysis as in (A) for the same dataset sorted manually. **C**: Cosine similarity between templates computed by averaging extra spikes identified by YASS (red) or KS (blue, top panels) or manual sorting (green, bottom panels) versus the template computed by averaging the “consensus” spikes identified by both spike-sorters. YASS extra spikes tend to be much more similar to the consensus template, consistent with the examples shown in Figure 17. In the left panels we scatterplot these cosine similarities against the PTP size of the unit; in the right panels against the firing rate of the extra identified spikes. Note that in many cases YASS identifies a large number of extra spikes that closely match the consensus template. **D**. Same analysis as in (C) for another dataset.

Figure 19 analyzes the non-consensus units, i.e., units that are found by YASS but not KS2, or vice versa. Overall we find that KS2-only units tend to have few spikes (in some cases so few spikes that it was challenging to estimate reliable templates) that tend to be more noisy, with a higher overall rate of outliers. The YASS-only units, on the other hand, tend to have lower PTP values but lower outlier scores in general.

**Figure 19:**
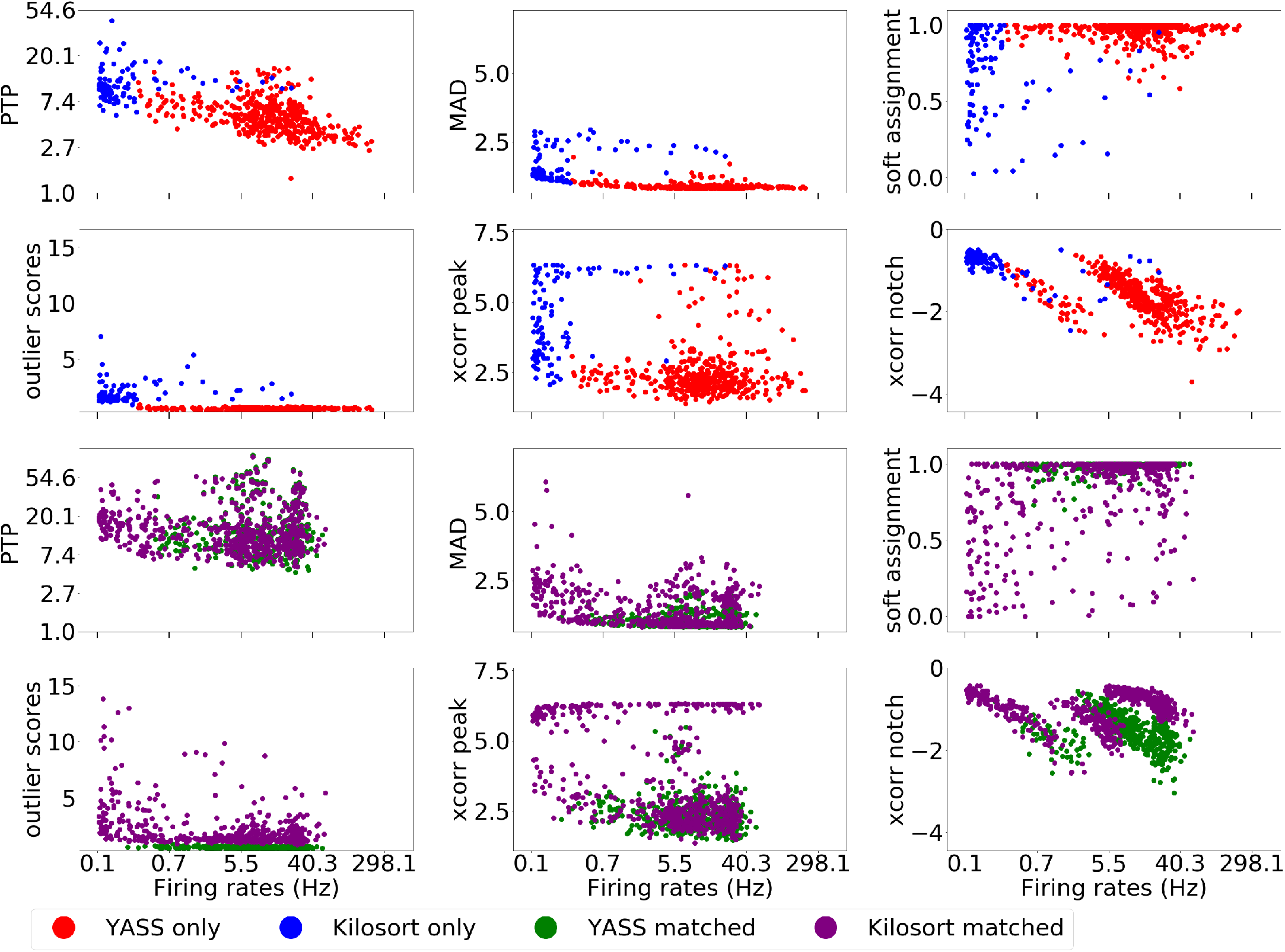
Comparing consensus and non-consensus units from YASS and Kilosort. Among units with firing rates bigger than 0.1 Hz, we found 444 unmatched YASS-only and 101 Kilosort-only units, compared to 491 (YASS) and 595 (Kilosort) matching consensus units (note that this is not a pairwise matching, due to oversplits). The top panels show non-consensus units, and the bottom panels show consensus units. We scatterplot the firing rate of each unit against: the unit’s PTP; the maximum absolute deviation (MAD) of the aligned cleaned spikes (overly large values indicate corruption of these spikes, due to e.g. deconvolution errors or over-merged units); the average soft-assignment probability (i.e., one minus the probability that a spike assigned to unit *i* should have been assigned to some other unit *j*, averaged over all spikes from unit *i*; small values indicate closely overlapping clusters that may correspond to over-splitting); the “outlier score” defined as the average norm of the whitened post-deconvolution residual around each spike (again, large values indicate corruption of these spikes); the maximal normalized cross-correlation (zero lag, one ms resolution) between unit *i* and all other units *j* (overly large values can indicate “fragmented” units, where for example axonal channels have been split from the somatic channels); and the minimal cross-correlation, defined similarly (a large “notch” in the cross-correlation function often indicates oversplitting, since if a cell is oversplit into two units *i* and *j* then *i* and *j* will tend not to spike at the same time).

Figure 20 compares the residuals computed by YASS and Kilosort (i.e., the difference between the raw data and the sum of the templates subtracted away during deconvolution by each of these methods). We see that Kilosort fails to subtract away many clearly visible spikes, while there are many fewer of these clear “missed spikes” visible in the YASS residual. Overall the YASS residual has a scale (as estimated by the MAD) of about 80% of that of the original “noise” level — i.e., YASS is able to explain a significant fraction of the “noise” by subtracting away many small contributions of distant spikes that are large and clearly identifiable on at least one channel. KS2, on the other hand, does not reduce the residual scale significantly.

**Figure 20:**
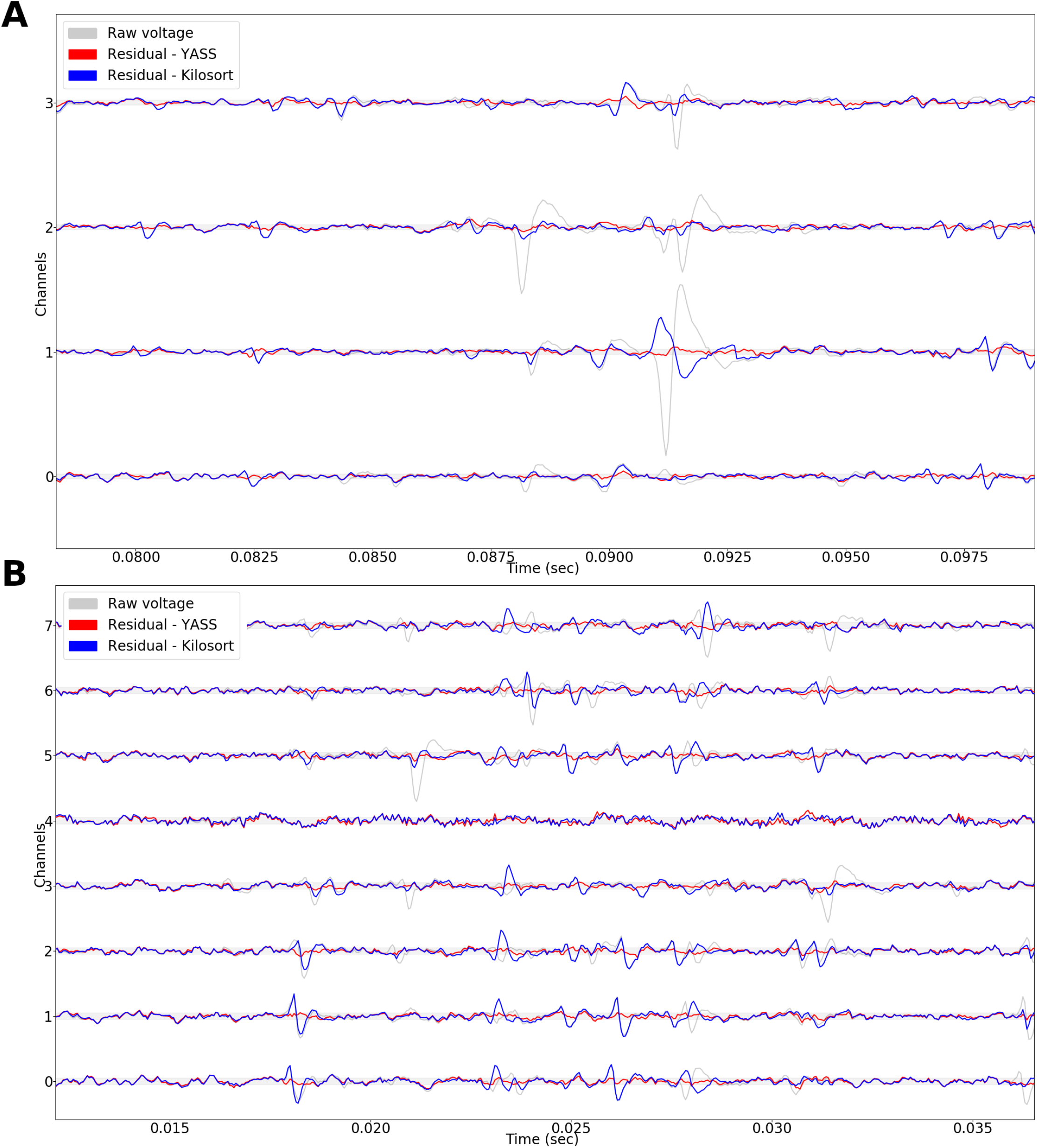
Residual analysis. **A**. A 20 ms example of YASS and Kilosort residuals on 5 channels. For large units, Kilosort occasionally leaves large residuals behind (c.f. Figure 8). **B**. Another example on 8 channels reveals some spike mis-assignment by Kilosort.

We additionally tested the full YASS pipeline on a number of “hybrid” datasets (following Rossant et al. (2015)), in which 20 neurons were added to an existing physiological recording containing 100-200 real neurons. YASS performed well on these datasets; here we restrict our discussion to a challenging dataset that contained 20 injected low-amplitude neurons (see Figure 21; on this dataset Kilosort did not detect any of the injected neurons at the default threshold). YASS is able to recover most spikes in most of the injected units, but for injected units with low firing rate and/or template size, the error rate increases sharply. We believe there may be room to continue to improve spike sorting performance in this challenging regime.

**Figure 21:**
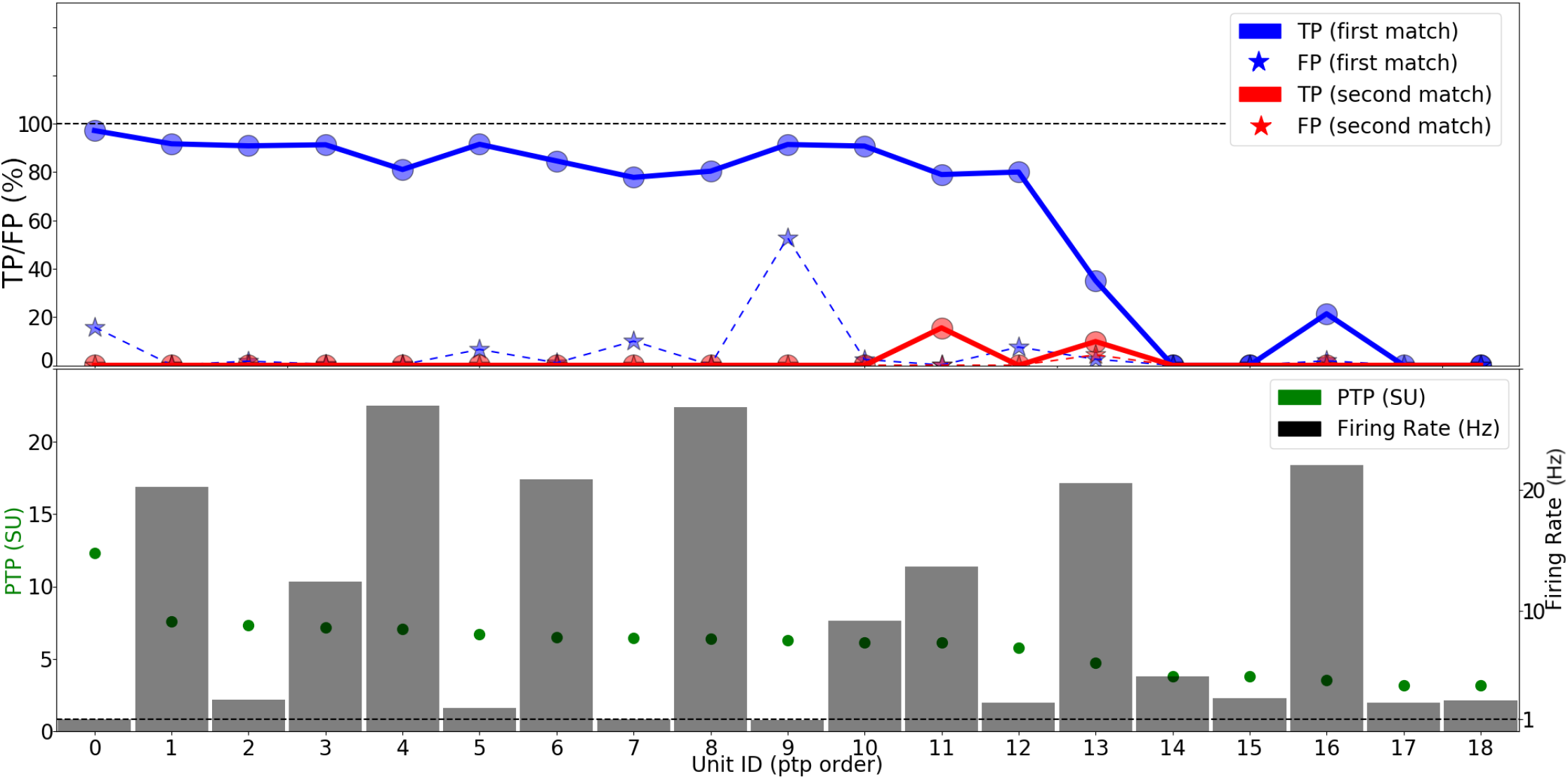
YASS results on hybrid semi-synthetic datasets containing low amplitude units. **Top**. The true positives (solid colors) and false positives (hatched colors) for the best (blue) and second best (red) matches for neurons sorted by yass (for clarity, a perfect match is a blue bar that contains 100% of the spikes with no other bars). **Bottom**. Firing rate and PTP amplitude for neurons shown above. (Note: Kilosort did not detect any of the injected units using the default thresholds).

### 3.3 Comparing decoding accuracy based on spike sorter output versus “unsorted” spikes

The results described in the previous subsection establish that YASS is able to extract units with receptive fields that are not found by human sorters or by Kilosort; moreover, YASS finds many spikes that are missed, and discards many outliers that are added, by human sorters or Kilosort. A natural question arises: how much of a difference do these extra units and spikes make when performing downstream analysis tasks with the resulting data? For example, do improvements in spike sorting lead to improvements in our ability to decode visual information from the retinal output?

To address this question, we analyzed retinal responses to flashed natural images. Specifically, static images from the ImageNet database (Deng et al., 2009) were flashed for 100 ms, with a gap of 400 ms of gray in between each flash. We then applied YASS and Kilosort2 to sort the retinal responses driven by these flashed stimuli. As a baseline inspired by Trautmann et al. (2019), we also extracted “unsorted” spikes by running the detect and de-duplication stage of the YASS pipeline and then assigning each spike to its primary channel (the channel on which the spike was largest) to obtain 512 unsorted “multi-units” (one multi-unit per electrode).

Then we trained simple linear decoders (Warland et al., 1997, Stanley et al., 1999) on these three sets of spike trains. We used a coarse temporal binning that captured both the onset and offset response driven by the flashed stimulus: the onset response was captured by summing spikes in the 30-150 ms window after image stimulus onset, and the offset 170-300 ms after image onset. We then appended the resulting binned spike counts from each unit into a vector summarizing the response to each image; therefore, the input to the decoder for each image was a 2*K* × 1 vector of binned spike counts, where *K* is the number of units extracted by each sorter. We used linear ridge regression to map this 2*K* × 1 input vector into an estimate of each pixel’s intensity, and then appended these intensity estimates to form the decoded image. There were a total of 10,000 non-repeated images in the dataset; we used a 9800-100-100 train-validation-test split to select the optimal ridge regression parameter for each of the three decoders and perform cross-validation. Finally, we quantified the error of the three decoders by computing the *R*^2^ between the true versus decoded images in the test dataset.

Figure 22 summarizes the results: we see that the decoder based on the YASS output significantly outperforms the Kilosort-based decoder, which in turn significantly outperforms the decoder based on unsorted spikes. Thus, improvements in spike sorting performance do translate to improvements in decoding accuracy in this case. We also experimented with a hybrid approach, in which we ran YASS, then ran the detection step on the residual (to collect spikes that might have been missed by YASS), then ran decoding using both the YASS units and the spikes detected in the pass through the residual. This hybrid approach did not improve the decoding accuracy, indicating that, as desired, YASS is extracting much of the visually-relevant information from the raw MEA output.

**Figure 22:**
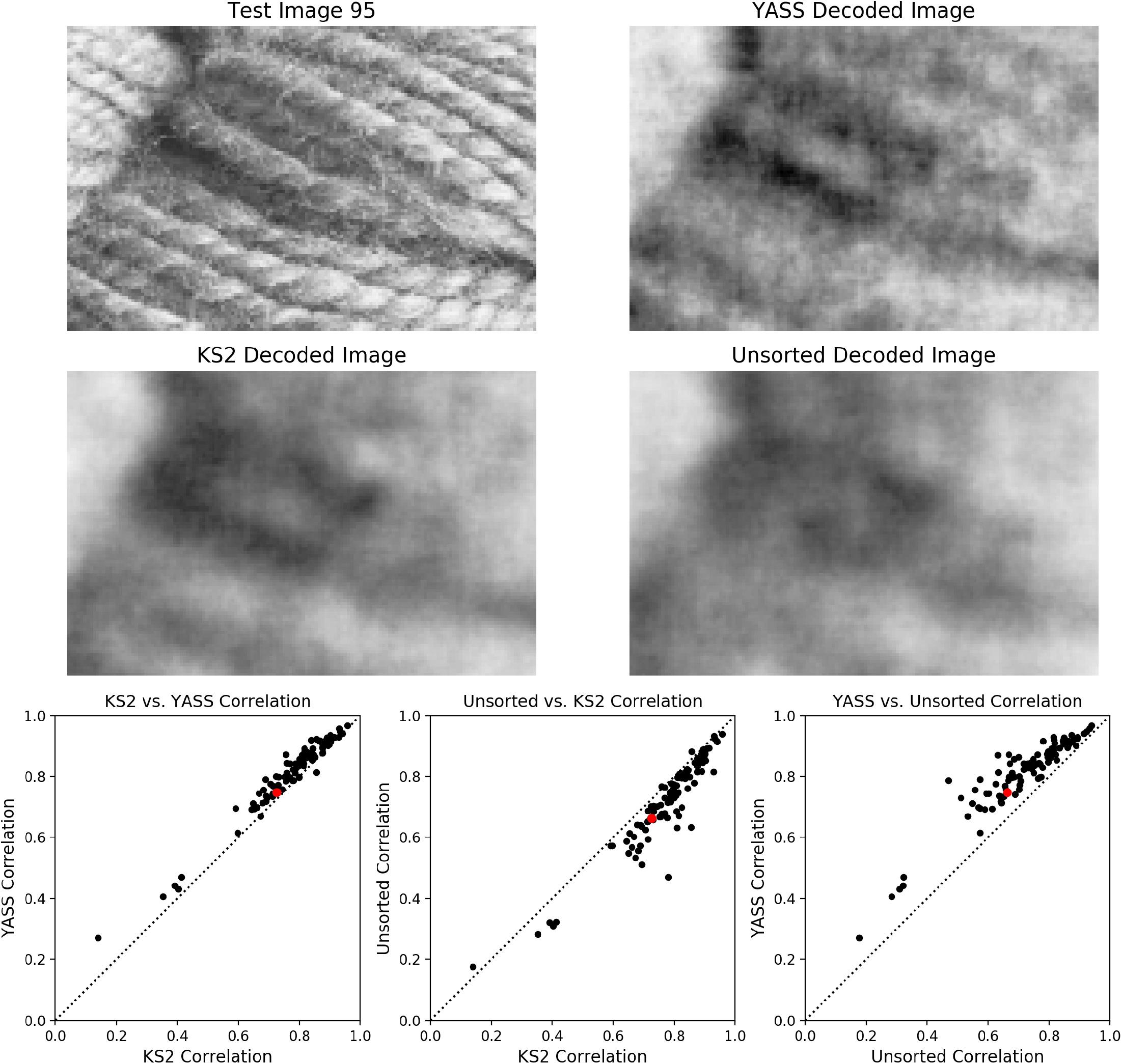
Comparing decoding accuracy based on YASS versus Kilosort output versus “unsorted” spikes. Top: example image from the test set, along with decoded images based on YASS output, Kilosort output, and unsorted spikes. Bottom: scatterplots comparing YASS versus Kilosort versus unsorted decoding accuracy. (Each dot corresponds to the correlation between the decoder and true image for one test image, so higher values are better; red dot indicates test image shown in top panels.) YASS leads to significantly more accurate decoding than Kilosort, and Kilosort leads to significantly more accurate decoding than unsorted spikes; i.e., even suboptimal spike sorting leads to better decoding than unsorted decoding.

### 3.4 Comparing sorts based on linear probe geometry versus the full planar MEA

The retinal ganglion layer is to a first approximation two-dimensional, as is the planar MEA used to perform the recordings analyzed here; thus, the recording device in this preparation is approximately matched to the geometry of the recording target. In contrast, this geometric matching usually does not hold for recordings in other parts of the brain, where typically we use probes with linear (approximately one-dimensional) geometry (Jun et al., 2017a) to target three-dimensional structures.

This raises a critical question: how much information is lost due to this mismatch between the recording device and neural target? Given the accurate sorting quality provided by YASS, we can address this question quantitatively by performing a simple experiment: first perform spike sorting given data obtained in a setting where the recording device and target geometry match (as we have already done here). Then perform spike sorting given data from a subset of electrodes chosen to emulate a linear probe geometry, and compare the results of these two sorts: e.g., quantify how many units are missed, how many mistaken splits or merges occur, and so on.

The results of this experiment are summarized in Figure 23. Two main conclusions can be drawn. First, we do see significant overmerging when comparing the units sorted from the “line” subset to the units sorted from the full array. While many line units match well with the corresponding array units (as quantified by “purity” — the proportion of spikes in the line unit that are also found in the best-matching array unit — and conversely “completeness” — the proportion of spikes in the array unit that are also found in the best-matching line unit), as seen in the dots that hug the 100% line in Figure 23, there are a number of line units that are merges of two or more array units; see Figure 24 for an example. In these cases, the limited context provided by the linear subset was insufficient to split two units that could be easily distinguished on channels that were distant from the line. Importantly, these overmerges could be found even in units with relatively large magnitude, i.e., PTP > 10.

**Figure 23:**
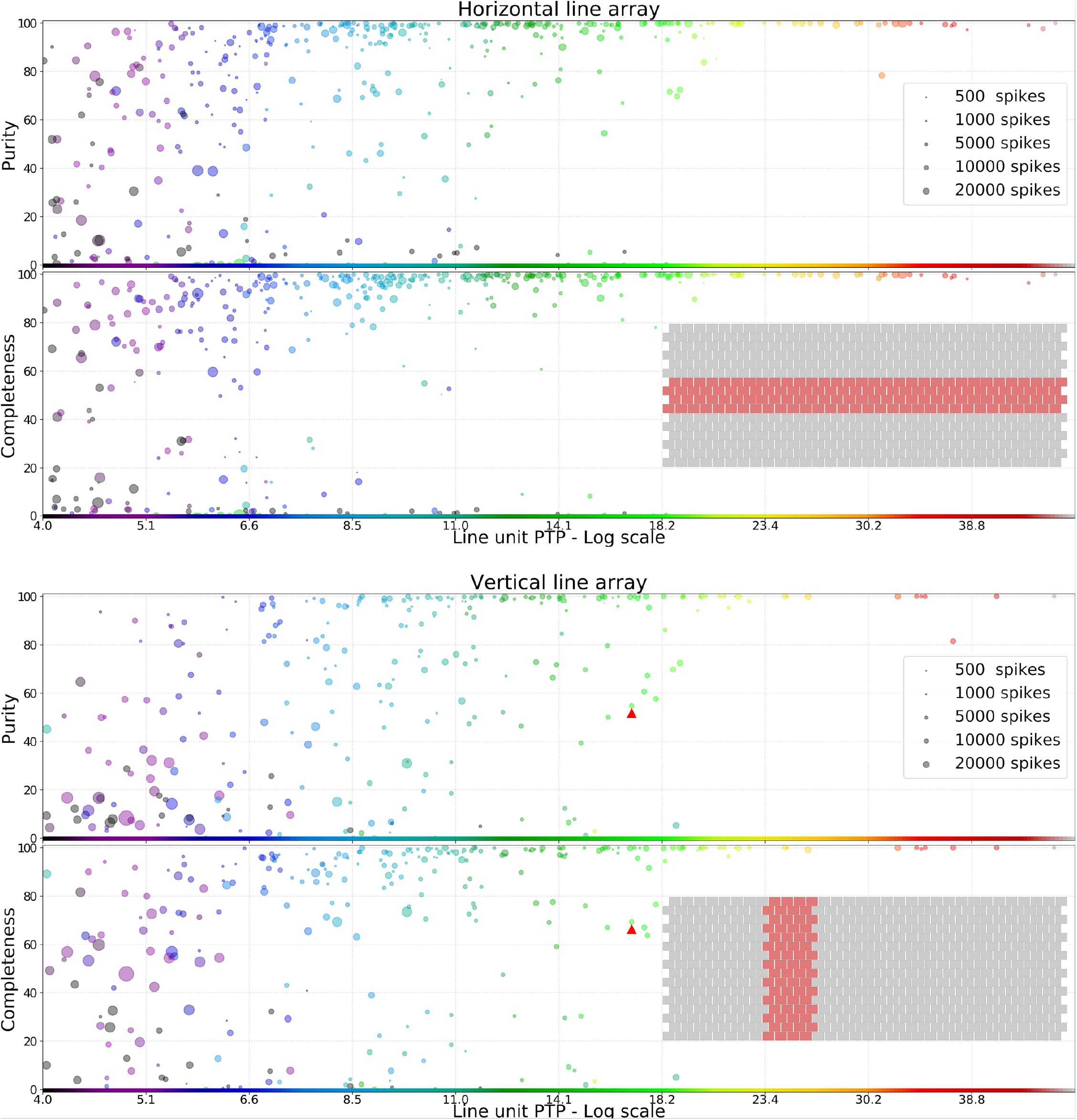
Comparing sorts based on linear probe geometry versus the full planar MEA. Purity and completeness (defined in the main text) shown for two linear probe geometries, one horizontal (top) and one vertical (bottom) with respect to the orientation of the full array. The geometry of the recording sites for the full array (gray boxes) and both linear probes (red boxes) are shown in the insets. X-axis indicates the maximum peak-to-peak (PTP) value of the line unit; y-axes the purity and completeness values from 0-100%. Each scatter circle represents a line unit. The size of the circles is linearly proportional to the number of spikes of that unit. The color of the circles represent the largest PTP of the best matched array units, with PTP measured on the linear channels, not the full array. The red triangle indicates the example overmerged line unit shown in Figure 24. Note that most high-PTP units have high purity and completeness scores, but there are exceptions (i.e., line units that do not match a single array unit well); on average, the purity and completeness scores tend to be higher for the horizontal than for the vertical linear geometry, because in this recording axons tend to have a more horizontal orientation.

**Figure 24:**
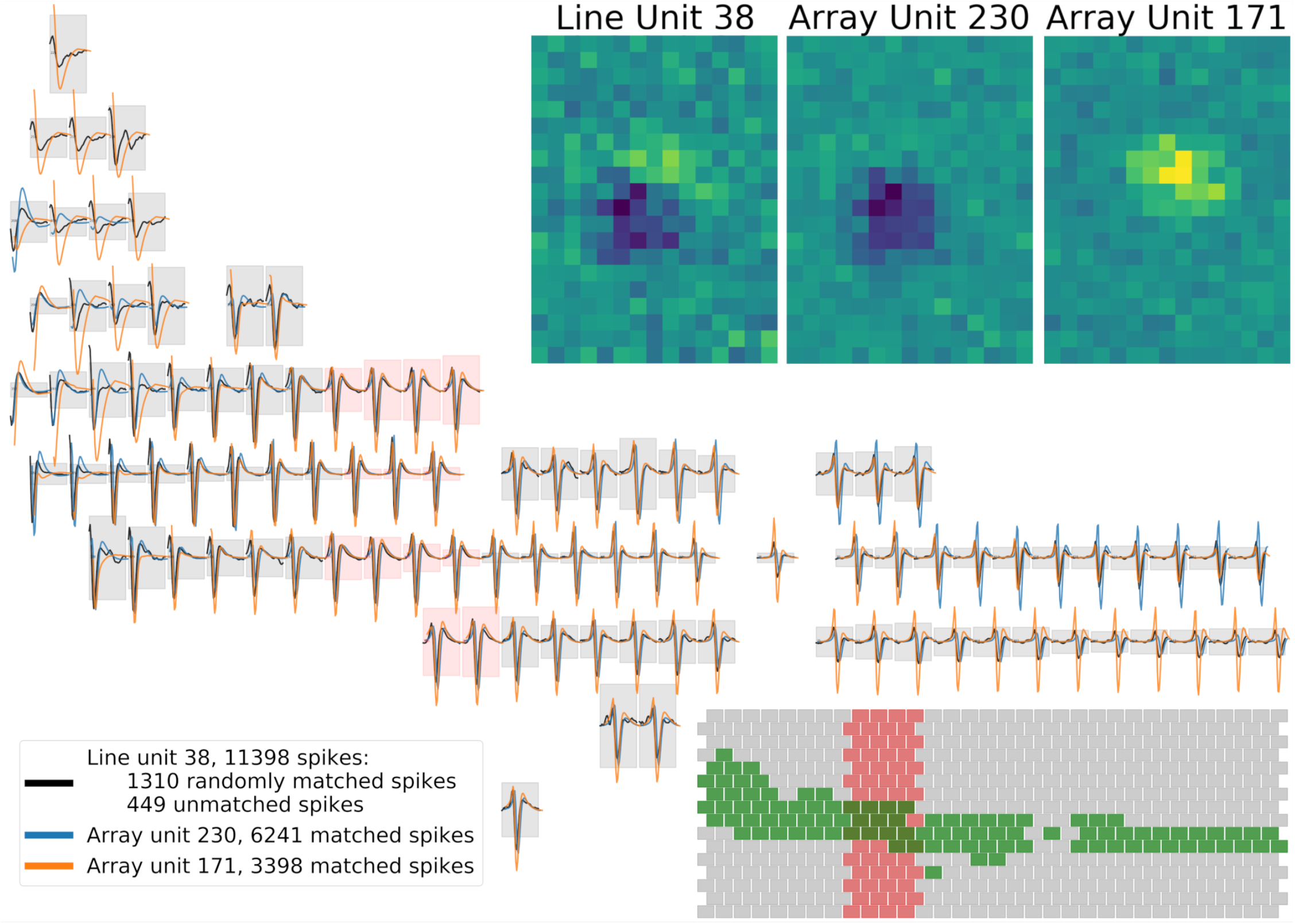
Example of a line unit that merges two array units. (This is the unit marked by a triangle in Figure 23.) The templates of the line and two array units are shown, plotting only channels that are active for at least one of the units. Lower right inset indicates the linear probe geometry (red boxes) and the active channels (green) in the context of the full array layout (gray). For each active channel, the red or gray box shown behind the templates marks ±1 standard voltage units (so a larger gray box indicates that the template on that channel is smaller). The cropped receptive fields (RFs) of all three units are shown in the upper right. On the linear probe channels (red boxes) the three templates are very similar, while outside on the full array, the templates of the two array units are significantly different. The line unit groups together spikes from these two array units (along with a smaller number of spikes that are not matched to any array unit, or are synchronous with several other apparently random array units), because on the linear probe channels it cannot differentiate between the two array units. This causes the line unit’s template and RF to be a weighted mean of the templates and RFs of the array units.

Second, the degree of overmerging depends strongly on the local geometry of the circuit relative to the linear probe geometry. In the dataset analyzed here, axons tended to be horizontally aligned. Therefore, a vertically-oriented linear probe had more trouble separating these axonal signals than a horizontally-oriented probe (as quantified with the relatively lower purity and completeness scores in the bottom plot of Figure 23 compared to the top), simply because horizontally-aligned axons will tend to contact fewer electrodes on the vertical probe, thus providing less information for the spike sorter to use to split neighboring units.

## 4 Conclusion and next steps

`YASS` achieves state-of-the-art, superhuman performance in a challenging dense multi-electrode spike sorting setting in which collisions and highly overlapping cell shapes are prevalent. This improved performance was enabled by a number of methodological innovations: neural network-based detection and denoising of spike waveforms, post-deconvolution “cleaning” of collided waveforms, triaging and iterative re-featurization for more robust clustering, and improved template estimation and deconvolution for better resolution of collided spikes. We also developed useful new visualization and diagnostic tools (Figures 17–20) that are applicable to compare any spike sorters.

We applied `YASS` to address two important basic neuroscientific questions: (1) does better spike sorting lead to better performance in downstream tasks such as decoding? (2) How much information is lost when we record from a quasi-linear probe compared to an MEA that better matches the geometry of the neural substrate? Regarding (1), we find that better spike sorting does indeed lead to better decoding performance, and even suboptimal spike sorting improves over decoding from “unsorted” spikes. This finding is consistent with the results of Todorova et al., but differs from the message of Trautmann et al. (2019), perhaps because the images we decode here have dimensionality that is orders of magnitude larger than the motor signals decoded in Trautmann et al. (2019). Regarding (2), we find that recording from a linear probe geometry can lead to a significant number of overmerges (Figures 23–24), and that the degree of overmerging depends strongly on the underlying spatial orientation of the targeted cells. Thus it is fair to conclude that some caution is warranted in interpreting spike trains extracted from linear probes, particularly when recording from circuits in which the targeted cells are aligned orthogonally to the probe.

We have focused here on MEA data collected from the primate retina. As noted above, this preparation is a useful spike sorting testbed for a number of reasons: the approximately two-dimensional substrate of the retinal ganglion layer matches the two-dimensional MEA here (enabling the quantification of linear probe accuracy discussed in section 3.4); the mosaic RF structure in the retina provides useful side information for scoring different spike sorting pipelines (c.f. Figure 16); and collision rates in the primate retina are high (Figure 13), presenting a useful challenge to available spike deconvolution algorithms. In our future work, we plan to explore the generality of the tools developed here, by pursuing applications to MEA data recorded in other parts of the nervous system. We expect that some of these applications will require extensions of the tools developed here; for example, tracking rapid template changes in low-firing-rate cells remains a challenge, as does deconvolution of very densely collided data, automatic detection of behavior-induced artifacts, and tracking drifting cells in chronic recordings over days (instead of the hours-long datasets analyzed here). We hope to address some of these challenges in the near future.

## Acknowledgements

We thank Gaute Einevoll, Espen Hagen, Ari Pakman, Calvin Tong, Yueqi Wang, and Weichi Yao for many helpful conversations. We also thank the developers of Kilosort/Kilosort2, JRClust/Ironclust, Mountainsort, Spyking Circus, and Herding Spikes for generously sharing their code. This work was partially supported by the Simons Foundation, the ONR, and NSF grants IIS-1546296 and NSF IIS-1430239. This research was also developed with funding from the Defense Advanced Research Project Agency (DARPA), Contract No. N66001-17-C-4002. The views, opinions and/or findings expressed are those of the author and should not be interpreted as representing the official views or policy of the Department of Defense of the U.S. Government.

## Appendix

### A Notation

We use the following conventions: scalars are lowercase italicized letters, e.g. *x*, constants such as max indices are represented by uppercase italicized letters, e.g. *N*, vectors are bolded lowercase letters, e.g. **x**, and matrices are bolded uppercase letters, e.g. **X**. Major notations used in the paper are summarized in Table 1.

**Table 1:** Summary table of notation used within the manuscript.

| Notation | Explanation | Default value (if exists) |
| --- | --- | --- |
| **Data Constants** |  |  |
| $T$ | Recording length (in timesteps) | |
| $C$ | Total number of channels | |
| $N$ | Number of detected spikes | |
| $C_{neigh}$ | Number of neighboring channels (including the main channel) | |
| **Parameters** |  |  |
| $R$ | Temporal window size of spike | 5 ms |
| $R_{NN}$ | Temporal window size of spike used for NN detection and denoising | 3 ms |
| $\tau$ | Threshold for spike detection under NN output | 0.5 |
| **Data structures** |  |  |
| $\mathbf{V} \in \mathbb{R}^{T \times C}$ | Recording | |
| $\mathbf{P} \in (0, 1)^{T \times C}$ | Sigmoid output of Neural Network Detection | |
| $\mathbf{X}, (\mathbf{X}_n)$ | Waveform (of spike $n$) | |
| $\mathbf{W}_k$ | Template of unit $k$ | |
| **Notations for MFM** |  |  |
| $K$ | Number of components in the generative model | |
| $\pi = (\pi_1 \dots \pi_k)$ | Component weights in the generative model | |
| $\mu_k, \mathbf{\Lambda}_k$ | Mean and precision matrix of component $k$ | |
| $\mathbf{z} = (z_1, \dots, z_n)$ | Component membership | |
| $\mu_0, \lambda_0, \mathbf{V}_0, \nu_0$ | Prior parameters for the Normal-Whishart | $\mathbf{0}, 0.01, \frac{1}{\nu_0} \mathbf{I}_D, D + 2$<br>$D$: # of feature dimensions |
| $\alpha$ | Prior concentration parameters for Dirichlet dist. | 1 |
| $\beta$ | Prior parameter for Poisson Distribution | 1 |
| $\hat{K}$ | Variational parameter for the point mass | |
| $\hat{\alpha}_k$ for $k = 1 : \hat{K}$ | Variational concentration parameters for the posterior Dirichlet Distribution | |
| $\hat{\mu}_k, \hat{\lambda}_k, \hat{\mathbf{V}}_k, \hat{\nu}_k$ for $k = 1 : \hat{K}$ | Variational parameters for the posterior Normal-Whishart | |

### B Additional details on the detection algorithm

#### B.1 Neural network training data

See Figure 25 for an overview of how we construct each training sample for the detection neural network.

**Figure 25:**
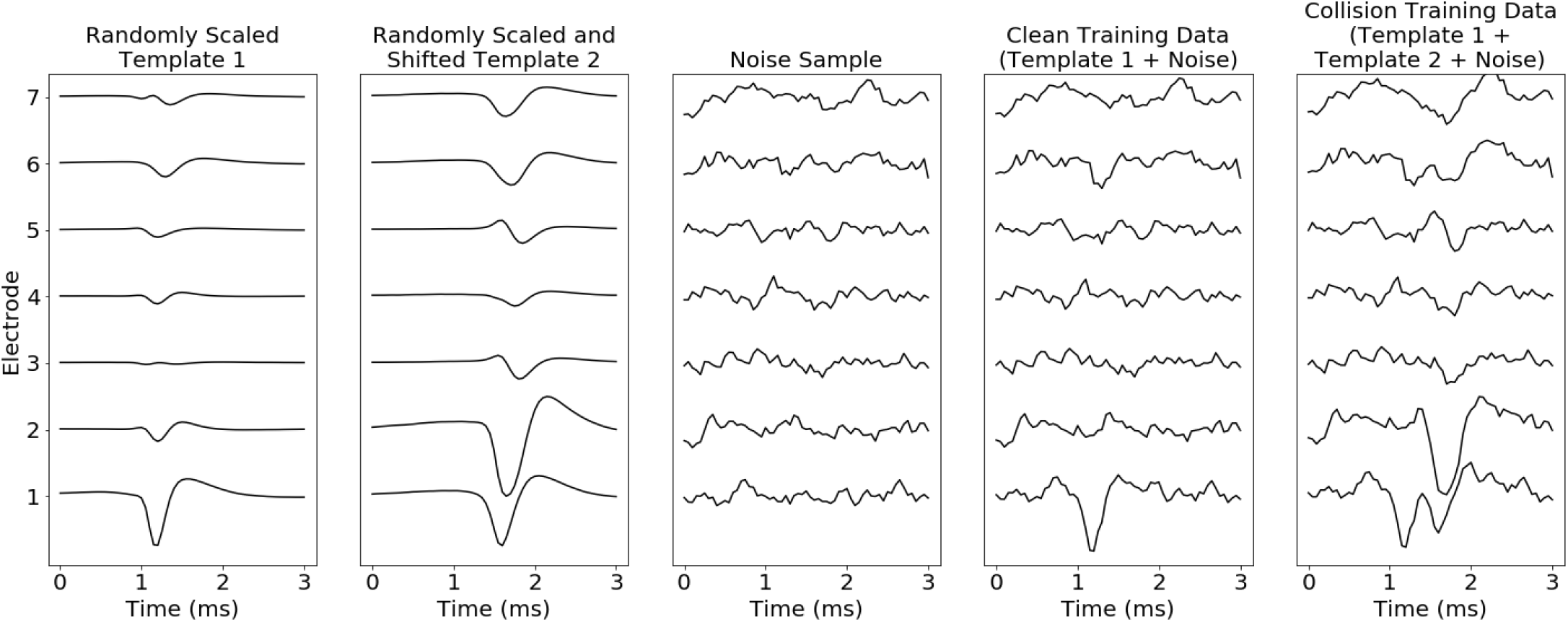
Illustration of Construction of NN Training Data. (Column 1) One template is randomly chosen from a previous sort and randomly scaled. This template is centered temporally. (Column 2) Another template may be randomly chosen, randomly scaled, and randomly shifted. (Column 3) A Gaussian noise sample (both temporally and spatially correlated) is drawn. (Column 4) A single spike sample is created by superimposing template 1 and noise. (Column 5) A collision sample is constructed by adding template 1, template 2, and noise. No more than 2 templates are used for creating collision samples. (Column 6) A misaligned sample is created by adding noise to a randomly shifted template. When the convolutional neural network is applied to the recording, our goal is to output no duplicate detection of the same spike. Thus, during training, samples such as column 4 and 5 have label 1 and samples such as column 3 and 6 have label 0.

We start with a library of clean templates from previous sorts. (Since the detection network operates locally in space and time, we only need cleaned templates on local 7-electrode patches here: a center electrode and the six surrounding electrodes.) We choose a template randomly, scale it randomly, and center it, and then (optionally) corrupt this clean template with a collision from a different template (with a random scale and shift). (We found that it was not necessary to form collisions using >2 spikes.)

Next we add indepedent noise samples to these templates. For this we need to construct a distribution over noise samples of dimension *R_NN_* × *C_neigh_* = 61 × 7 here. Following previous work Pillow et al. we use a spatiotemporally separable, stationary Gaussian noise model. We find “noise” snippets in the raw data by excluding any spikes detected by simple thresholding with a low threshold (3 standardize units). We then rescale these snippets so that their standard deviation has unit standardized units and compute the temporal and spatial covariance matrix. We approximate the resulting temporal covariance with a Toeplitz (temporally stationary) matrix, and the spatial covariance with a rotation-symmetric matrix (to enforce spatial stationarity), then form the Kronecker product to obtain the full spatiotemporal covariance matrix.

Given an unlimited stream of samples from the above generative model, we train the neural network detector as a classifier that takes as input waveforms 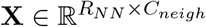, and returns a soft-max output between (0, 1) that estimates the presence of a spike. Single-spike and collision samples have positive labels; noise and off-centered samples have negative labels. We use an equal proportion of positive and negative labels, and the misaligned spike is added 70% of the time to reflect the high collision rate of retina recordings.

#### B.2 Neural network structure

The proposed architecture is the detector is shown in Figure 26. To build a faster network, temporal and spatial filters are separated. The first two layers extract temporal features and the third layer combines the features in spatially neighboring channels. The first layer applies a one-dimensional convolutional neural network using *K*_1_ filters of size *R_NN_*. The second layer maps *K*_1_ features to *K*_2_ features at each location. The third layer transforms *C_neigh_* × *K*_2_ features into a single number. Rectified linear unit nonlinearities (ReLU) are used as activation functions and the sigmoid transform is applied at the end to get a value in (0, 1). The network is learned by minimizing the cross-entropy loss using the Adam update rule (Kingma and Ba, 2015). For the retinal recordings analyzed here, *K*_1_ and *K*_2_ are set to 16 and 8. The learning rate of Adam is set to 0.0001.

**Figure 26:**
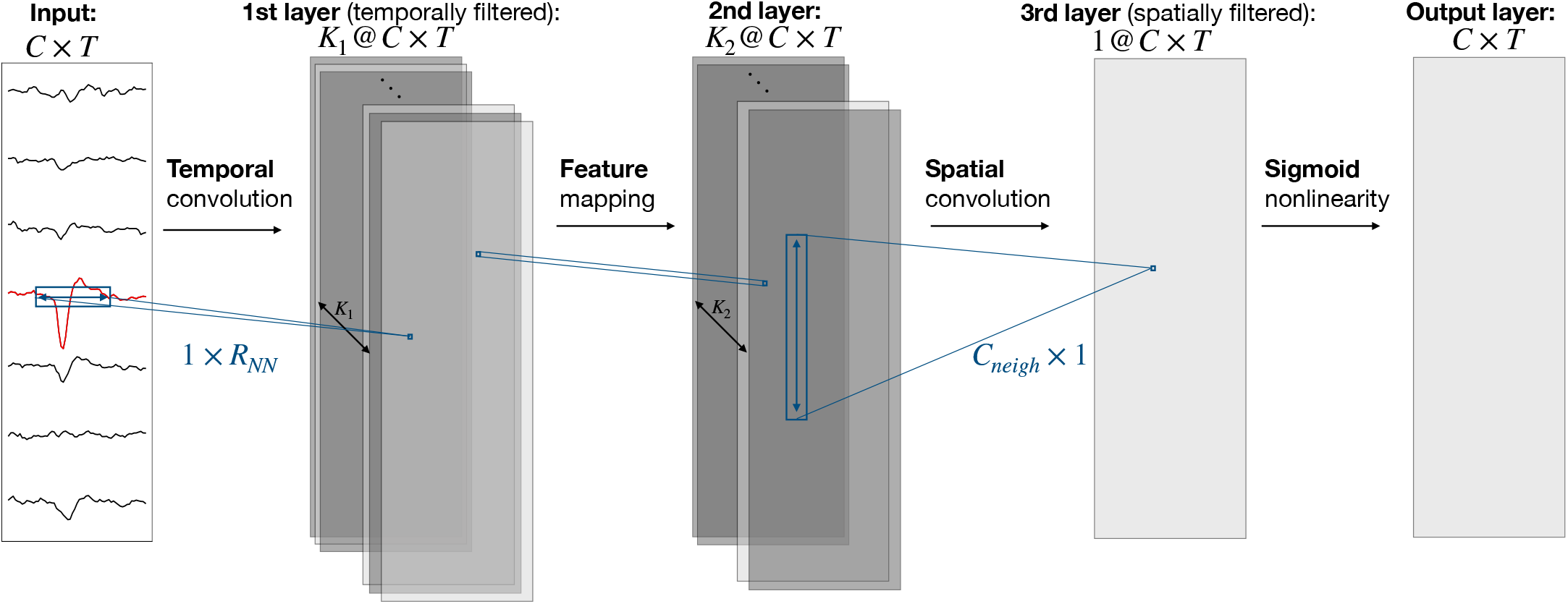
Visualization of the architecture of Neural Network detector. ReLUs are used as activation functions.

After training, given a recording, 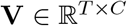, the neural network outputs an array, **P** ∈ (0, 1)^*T*×*C*^, where **P**_(*t*,*c*)_ is the output of the neural network (applied convolutionally in space and time) with input a temporal and spatial snippet of **V** around *t* and *c*. Given the output, **P**, temporal and spatial local maxima that cross a threshold, *τ*, are considered as detected events. As a result, the detector outputs pairs of spike times and corresponding channels, (*t*_1_, *c*_1_), …, (*t_N_*, *c_N_*).

### C Additional details on the denoising algorithm

The training data for the neural network denoiser is constructed as above for the neural network detector. Instead of the 0-1 label output by the detector, the denoiser output targets the ground truth template without any noise or collision superimposed.

We found that denoising on single channels already gave good performance. Thus the denoiser takes a waveform of size *R_NN_* as input. The network consists of three hidden layers with one-dimensional convolutional neural network and ReLU activation functions. At the end, a linear mapping is applied to output a vector of size *R_NN_*. The NN is trained to minimize the L2 distance between the output and the ground truth clean template. For the retina, the three hidden layers have 16, 8, and 4 filters of size 5, 11, 21 respectively. We use the same training parameters as described for the detector network above.

### D Additional details on the Mixture of Finite Mixture (MFM) model

In the clustering step we use a Mixture of Finite Mixture model (MFM) described in Miller and Harrison (2018). To estimate the posterior distribution we use a structured variational distribution (Hoffman and Blei, 2015) which preserves dependencies between parameters, leading to a more accurate posterior approximation than simpler unstructured variational approximations. Finally, we use split-merge steps based on Hughes and Sudderth to estimate a reasonable number of clusters.

#### D.1 Generative Model

We assume the following generative model (Miller and Harrison, 2018):

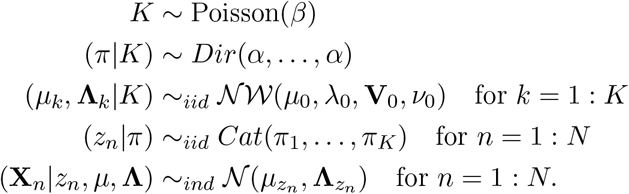

The joint distribution of *K*, *π*, *μ*, **Λ**, **z**, **X** is given by:

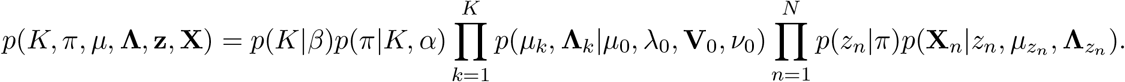

#### D.2 Inference model and estimation

The following structured variational distribution (Hoffman and Blei, 2015) is used for inference:

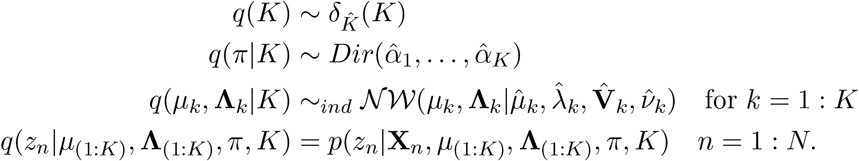

Also, the exact conditional is used for *q*(*z_n_*|*μ*_(1:*K*)_, **Λ**_(1:*K*)_, *π*, *K*), since it is tractable for the Gaussian mixture model. Given the global parameters, the conditional can be computed as:

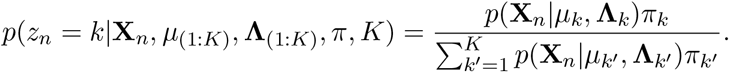

In summary, the variational joint distribution is then given by:

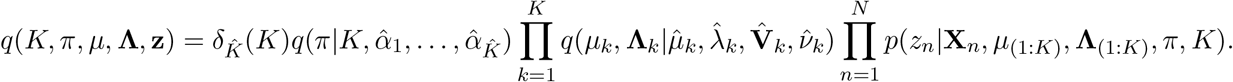

The variational parameters are estimated by maximizing the Evidence Lower Bound (ELBO):

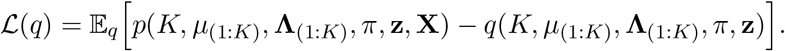

To estimate the variational parameters of *μ*_(1:*K*)_, **Λ**_(1:*K*)_, *π*, we use an approach similar to SSVI-A in Hoffman and Blei (2015). However, instead of stochastic updates, where *μ*_(1:*K*)_, **Λ**_(1:*K*)_, *π* are randomly sampled to estimate *q*(*z_n_*|*μ*_(1:*K*)_, **Λ**_(1:*K*)_, *π*, *K*), we use MAP estimation. This simplification is reasonable as the variational distributions are likely to have small variance due to large sample size. To infer the number of components, *K*, the exact birth and merge approach in Hughes and Sudderth is used.

### E Additional details on spline interpolation within deconvolution

As noted in the main text, subsample alignment errors between templates and the observed sampled data can lead to large residuals in the deconvolution step. In order to correct for these alignment error, we begin by estimating the temporal offset between the true versus sampled objective function peaks. To estimate the true peak locations, we fit a quadratic function to the samples around each detected peak and solve for the location where the maximum is obtained. With this information, the convolved templates can be shifted via interpolation prior to subtraction from the objective in order to align their peaks with the estimated peaks.

We use a Basis Spline (de Boor, 2001) representation of each convolved template. B-splines are capable of representing any spline function of a given order with sparsely supported basis functions whose support grows linearly with the order of the spline representation (i.e., the complexity of evaluation at a single location remains constant in the length of the templates). For example, a third order or “cubic” spline interpolation can be performed with the evaluation of only 4 basis functions at any given location, no matter the length of the function or number of knots. Therefore on the fly interpolation of a template represented as a combination of B-spline bases retains both the accuracy of standard spline interpolation and the linear computational complexity of linear interpolation. Moreover, evaluation of B-spline basis functions can be performed efficiently on the fly and lends itself to fast implementation on the GPU. We have implemented this functionality and in practice the cost incurred by performing interpolated subtraction is negligible relative to uncorrected subtraction.

## Footnotes

‡ DARPA Neural Engineering System Design program BAA-16-09

* On the other hand, our focus on the primate retina means that in this work we will de-emphasize some issues that are critical in different parts of the nervous system: for example, firing rate dependent spike adaptation, or drifts and nonstationarities in the size and shape of spikes on the MEA (both of which are present in the retina but pose more serious challenges in in vivo recordings). Moreover, issues such as “glitches” due to behavior are absent in our in vitro retinal recordings. However, we believe that the pipeline developed here can be adapted to address these issues, and we plan to pursue these directions in future work.

† In a previous version of this work (published in NIPS 2017), we trained a network to triage collision events. We have found that it is more data-efficient to denoise instead of triage these collisions if possible.

‡ A small technical note here; we do not denoise the distant channels, since the denoiser is trained on aligned spikes, and due to axonal propagation, spikes on distant channels may be poorly aligned. In the future, this issue could potentially be handled by training a denoising NN with more poorly-aligned training data, but we have not yet pursued this direction systematically.

§ We restrict attention to temporally isolated spikes here to avoid burst-dependent changes in spike height.

## Notes

https://github.com/paninski-lab/yass

## References

D Carlson, V Rao, J Vogelstein, and L Carin. Real-Time Inference for a Gamma Process Model of Neural Spiking. NIPS, 2013.

David E Carlson, Joshua T Vogelstein, Qisong Wu, Wenzhao Lian, Mingyuan Zhou, Colin R Stoetzner, Daryl Kipke, Douglas Weber, David B Dunson, and Lawrence Carin. Multichannel electrophysiological spike sorting via joint dictionary learning and mixture modeling. IEEE TBME, 61(1):41–54, 2014.

Jason E. Chung, Jeremy F. Magland, Alex H. Barnett, Vanessa M. Tolosa, Angela C. Tooker, Kye Y. Lee, Kedar G. Shah, Sarah H. Felix, Loren M. Frank, and Leslie F. Greengard. A fully automated approach to spike sorting. Neuron, 95:1381, 2017.

Carl de Boor. A Practical Guide to Splines. Springer-Verlag New York, 2001.

J. Deng, W. Dong, R. Socher, L.-J. Li, K. Li, and L. Fei-Fei. ImageNet: A Large-Scale Hierarchical Image Database. In CVPR09, 2009.

Chaitanya Ekanadham, Daniel Tranchina, and Eero P Simoncelli. A unified framework and method for automatic neural spike identification. J. Neuro. Methods, pages 47–55, jan. ISSN 1872-678X. doi: 10.1016/j.jneumeth.2013.10.001.

Greg D. Field, Alexander Sher, Jeffrey L. Gauthier, Martin Greschner, Jonathon Shlens, Alan M. Litke, and E. J. Chichilnisky. Spatial properties and functional organization of small bistratified ganglion cells in primate retina. 27(48):13261–13272, 2007. ISSN 0270-6474. doi: 10.1523/JNEUROSCI.3437-07.2007.

Felix Franke, Michal Natora, Clemens Boucsein, Matthias H J Munk, and Klaus Obermayer. An online spike detection and spike classification algorithm capable of instantaneous resolution of overlapping spikes. J. Comp. Neuro., (1-2):127–148, aug. ISSN 1573-6873. doi: 10.1007/s10827-009-0163-5.

John A Hartigan, Pamela M Hartigan, et al. The dip test of unimodality. The annals of Statistics, 13(1):70–84, 1985.

Gerrit Hilgen, Martino Sorbaro, Sahar Pirmoradian, Jens-Oliver Muthmann, Ibolya Kepiro, Simona Ullo, Cesar Juarez Ramirez, Alessandro Maccione, Luca Berdondini, Vittorio Murino, et al. Unsupervised spike sorting for large scale, high density multielectrode arrays. Cell Reports, 18(10):2521–2532, 2017.

Matthew Hoffman and David Blei. Stochastic structured variational inference. In Artificial Intelligence and Statistics, pages 361–369, 2015.

Michael C. Hughes and Erik Sudderth. Memoized Online Variational Inference for Dirichlet Process Mixture Models. In NIPS, pages 1133–1141.

James J Jun, Nicholas A Steinmetz, Joshua H Siegle, Daniel J Denman, Marius Bauza, Brian Barbarits, Albert K Lee, Costas A Anastassiou, Alexandru Andrei, Çağatay Aydin, et al. Fully integrated silicon probes for high-density recording of neural activity. Nature, 551(7679):232, 2017a.

James Jaeyoon Jun, Catalin Mitelut, Chongxi Lai, Sergey Gratiy, Costas Anastassiou, and Timothy D Harris. Real-time spike sorting platform for high-density extracellular probes with ground-truth validation and drift correction. bioRxiv, page 101030, 2017b.

Shabnam N Kadir, Dan F M Goodman, and Kenneth D Harris. High-dimensional cluster analysis with the masked EM algorithm. Neural computation, (11):2379–94, nov. ISSN 1530-888X. doi: 10.1162/NECO_a_00661.

Diederik Kingma and Jimmy Ba. Adam: A method for stochastic optimization. ICLR, 2015.

Edwin M Knox and Raymond T Ng. Algorithms for mining distance-based outliers in large datasets. In VLDB, pages 392–403. Citeseer, 1998.

Jeffrey W Miller and Matthew T Harrison. Mixture models with a prior on the number of components. Journal of the American Statistical Association, 2018.

Jens-Oliver Muthmann, Hayder Amin, Evelyne Sernagor, Alessandro Maccione, Dagmara Panas, Luca Berdondini, Upinder S. Bhalla, and Matthias H. Hennig. Spike detection for large neural populations using high density multielectrode arrays. Frontiers in Neuroinformatics, 9:28, 2015.

Marius Pachitariu. kilosort2, 2019.

Marius Pachitariu, Nicholas A Steinmetz, Shabnam N Kadir, Matteo Carandini, and Kenneth D Harris. Fast and accurate spike sorting of high-channel count probes with kilosort. In NIPS, pages 4448–4456, 2016.

Nikhil Parthasarathy, Eleanor Batty, William Falcon, Thomas Rutten, Mohit Rajpal, EJ Chichilnisky, and Liam Paninski. Neural networks for efficient bayesian decoding of natural images from retinal neurons. In Advances in Neural Information Processing Systems, pages 6434–6445, 2017.

Jonathan W Pillow, Jonathon Shlens, E J Chichilnisky, and Eero P Simoncelli. A model-based spike sorting algorithm for removing correlation artifacts in multi-neuron recordings. PloS one, (5):e62123, jan. ISSN 1932-6203. doi: 10.1371/journal.pone.0062123.

Cyrille Rossant, Shabnam N Kadir, Dan FM Goodman, John Schulman, Mariano Belluscio, Gyorgy Buzsaki, and Kenneth D Harris. Spike sorting for large, dense electrode arrays. bioRxiv, page 015198, 2015.

G. Stanley, F. Li, and Y. Dan. Reconstruction of natural scenes from ensemble responses in the lateral geniculate nucleus. Journal of Neuroscience, 95(18):8036–8042, 1999.

Ruoxi Sun and Liam Paninski. Scalable approximate Bayesian inference for particle tracking data. In Proceedings of the 35th International Conference on Machine Learning, 2018.

Nicholas Swindale and Martin Spacek. Spike sorting for polytrodes: a divide and conquer approach. Frontiers in Systems Neuroscience, 8:6, 2014.

Sonia Todorova, Patrick Sadtler, Aaron Batista, Steven Chase, and Valérie Ventura. To sort or not to sort: the impact of spike-sorting on neural decoding performance. J. Neural Eng., (5):056005, aug. ISSN 1741-2552. doi: 10.1088/1741-2560/11/5/056005.

EM Trautmann, SD Stavisky, Lahiri S, Ames KC, Kaufman MT, O’Shea DJ, Vyas S, Sun X, Ryu SI, Ganguli S, and Shenoy KV. Accurate estimation of neural population dynamics without spike sorting. Neuron, 103(2): 292–308, 2019.

D. Warland, P. Reinagel, and M. Meister. Decoding visual information from a population of retinal ganglion cells. Journal of Neurophysiology, 78(5):2336–2350, 1997.

Martin Weigert, Uwe Schmidt, Tobias Boothe, Andreas Müller, Alexandr Dibrov, Akanksha Jain, Benjamin Wilhelm, Deborah Schmidt, Coleman Broaddus, Siân Culley, et al. Content-aware image restoration: pushing the limits of fluorescence microscopy. Nature methods, 15(12):1090, 2018.

Pierre Yger, Giulia LB Spampinato, Elric Esposito, Baptiste Lefebvre, Stéphane Deny, Christophe Gardella, Marcel Stimberg, Florian Jetter, Guenther Zeck, Serge Picaud, Jens Duebel, and Olivier Marre. A spike sorting toolbox for up to thousands of electrodes validated with ground truth recordings in vitro and in vivo. 7:e34518, mar 2018. doi: 10.7554/eLife.34518.

Young-Gyu Yoon, Peilun Dai, Jeremy Wohlwend, Jae-Byum Chang, Adam H Marblestone, and Edward S Boyden. Feasibility of 3d reconstruction of neural morphology using expansion microscopy and barcode-guided agglomeration. Frontiers in computational neuroscience, 11:97, 2017.

Tong Zhang. Adaptive forward-backward greedy algorithm for learning sparse representations. IEEE transactions on information theory, 57(7):4689–4708, 2011.

